# The evolutionary history of Nebraska deer mice: local adaptation in the face of strong gene flow

**DOI:** 10.1101/152694

**Authors:** Susanne P. Pfeifer, Stefan Laurent, Vitor C. Sousa, Catherine R. Linnen, Matthieu Foll, Laurent Excoffier, Hopi E. Hoekstra, Jeffrey D. Jensen

## Abstract

The interplay of gene flow, genetic drift, and local selective pressure is a dynamic process that has been well studied from a theoretical perspective over the last century. Wright and Haldane laid the foundation for expectations under an island-continent model, demonstrating that an island-specific beneficial allele may be maintained locally if the selection coefficient is larger than the rate of migration of the ancestral allele from the continent. Subsequent extensions of this model have provided considerably more insight. Yet, connecting theoretical results with empirical data has proven challenging, owing to a lack of information on the relationship between genotype, phenotype, and fitness. Here, we examine the demographic and selective history of deer mice in and around the Nebraska Sand Hills, a system in which variation at the *Agouti* locus affects cryptic coloration that in turn affects the survival of mice in their local habitat. We first genotyped 250 individuals from eleven sites along a transect spanning the Sand Hills at 660,000 SNPs across the genome. Using these genomic data, we found that deer mice first colonized the Sand Hills following the last glacial period. Subsequent high rates of gene flow have served to homogenize the majority of the genome between populations on and off the Sand Hills, with the exception of the *Agouti* pigmentation locus. Furthermore, mutations at this locus are strongly associated with the pigment traits that are strongly correlated with local soil coloration and thus responsible for cryptic coloration.

## INTRODUCTION

Characterizing the conditions under which a population may adapt to a new environment, despite ongoing gene flow with its ancestral population, remains a question of central importance in population and ecological genetics. Hence, many studies have attempted to characterize selection-migration dynamics. Early studies focused on the distributions of phenotypes using ecological data to estimate migration rates (*e.g.*, Clarke & Murray 1962; Endler 1977; Cook & Mani 1980); whereas later studies, focused primarily on Mendelian traits, documented patterns of genetic variation at a causal locus and contrasted that with putatively neutral loci to estimate selection and migration strengths (*e.g.*, Ross & Keller 1995; Hoekstra *et al*. 2004; Stinchcombe *et al*. 2004, and see review of Linnen *et al*. 2010). Only recently have such studies expanded to examine whole genome responses (see Jones *et al*. 2012; Feulner *et al*. 2015).

These selection-migration dynamics are well grounded in theory. Haldane (1930) first noted that migration may disrupt the adaptive process if selection is not sufficiently strong to maintain a locally beneficial allele. Bulmer (1972) went on to describe a two-alleles two-demes model in which such a loss is expected to occur if *m/s* > *a*/(1-*a*), where *m* is the rate of migration, s is the selection coefficient in population 1, and *a* is the ratio of selection coefficients between populations. More recent related work suggests that in order for a beneficial allele to reach fixation, *s* must be much larger than *m* (*e.g.*, Lenormand 2002; Yeaman & Otto 2011, and see review of Tigano & Friesen 2016).

In addition to fixation probabilities, the migration load induced by the influx of locally deleterious mutations entering the population has also been well studied. Haldane (1957) found that the number of selective deaths necessary to maintain differences between populations is proportional to the number of locally maladapted alleles migrating into the population. Thus, the fewer loci necessary for the diverging locally adaptive phenotype, the lower the resulting migration load. More recently, Yeaman & Whitlock (2011) argued that with gene flow, the genetic architecture underlying an adaptive phenotype is expected to have fewer and larger-effect alleles compared to neutral expectations under models without migration (*i.e.*, an exponential distribution of effect sizes (and see Rafajlovic *et al*. 2017)). As recombination between these locally beneficial alleles may result in maladapted intermediate phenotypes, this model further predicts a genomic clustering of the underlying mutations (Maynard Smith 1977; Lenormand & Otto 2000). The relative advantage of this linkage will increase as the population approaches the migration threshold, at which local adaptation becomes impossible (Kirkpatrick & Barton 2006). Taken together, these studies make a number of testable predictions relating migration rate with the expected number of sites underlying the locally beneficial phenotype at the locus in question, the average effect size of the alleles, the clustering of beneficial mutations, and the conditions under which a locally beneficial allele may be lost, maintained, or fixed in a population.

These theoretical expectations, however, have proven difficult to evaluate in natural populations owing to a dearth of systems with concrete links between genotype, phenotype, and fitness. In one well-studied system, deer mice (*Peromyscus maniculatus)* inhabiting the light-colored Sand Hills of Nebraska (formed within the last 8000 years following the end of the Wisconsin glacial period (Loope & Swinehart 2000)) have evolved lighter coloration than conspecifics on the surrounding dark soil (Dice 1941; Dice 1947). Using a combination of laboratory crosses and hitchhiking-mapping, the *Agouti* gene, which encodes a signaling protein known to play a key role in mammalian pigment-type switching and color patterning (Jackson 1994; Mills & Patterson 1994; Barsh 1996), and see review of Manceau *et al*. 2010), has been implicated in adaptive color variation (Linnen *et al*. 2009). Moreover, using association mapping, specific candidate mutations have been linked to variation in specific pigment traits, which in turn contribute to differential survival from visually hunting avian predators (Linnen *et al*. 2013).

While previous research in this system has established links between genotype, phenotype, and fitness, this work has focused on a single ecotonal population located near the northern edge of the Sand Hills. Thus, the dynamic interplay of migration, selection, and genetic drift, as well as the extent of genotypic and phenotypic structuring among populations, remains unknown. To address these questions, we have sampled hundreds of individuals across a transect spanning the Sand Hills and neighboring populations to the north and south. Combining extensive per-locale soil color measurements, per-individual phenotyping, and 660,000 SNPs genome-wide, this dataset provides a unique opportunity to evaluate theoretical predictions of local adaptation with gene flow on a genomic scale.

## RESULTS & DISCUSSION

### Environmental and phenotypic variation in and around the Nebraska Sand Hills

We collected *Peromyscus maniculatus luteus* individuals (*N* = 266) as well as soil samples (*N* = 271) from 11 locations spanning a 330 km transect across the Sand Hills of Nebraska (Figure 1; Supplementary Table 1). As expected, we found that soil color differed significantly between “on” Sand Hills and “off” Sand Hills locations (*F_1_* = 307.4, *P* < 1 x 10^−15^; Figure 2). Similarly, five of the six mouse color traits also differed significantly between the two habitats (Figure 2). In contrast, mice trapped “on” versus “off’ of the Sand Hills did not differ in other non-pigmentation traits including total body length, tail length, hind foot length, or ear length (see Supplementary Materials). Together, these results are consistent with strong divergent selection between habitat types on multiple color traits, but not other morphological traits.

**Figure 1.**
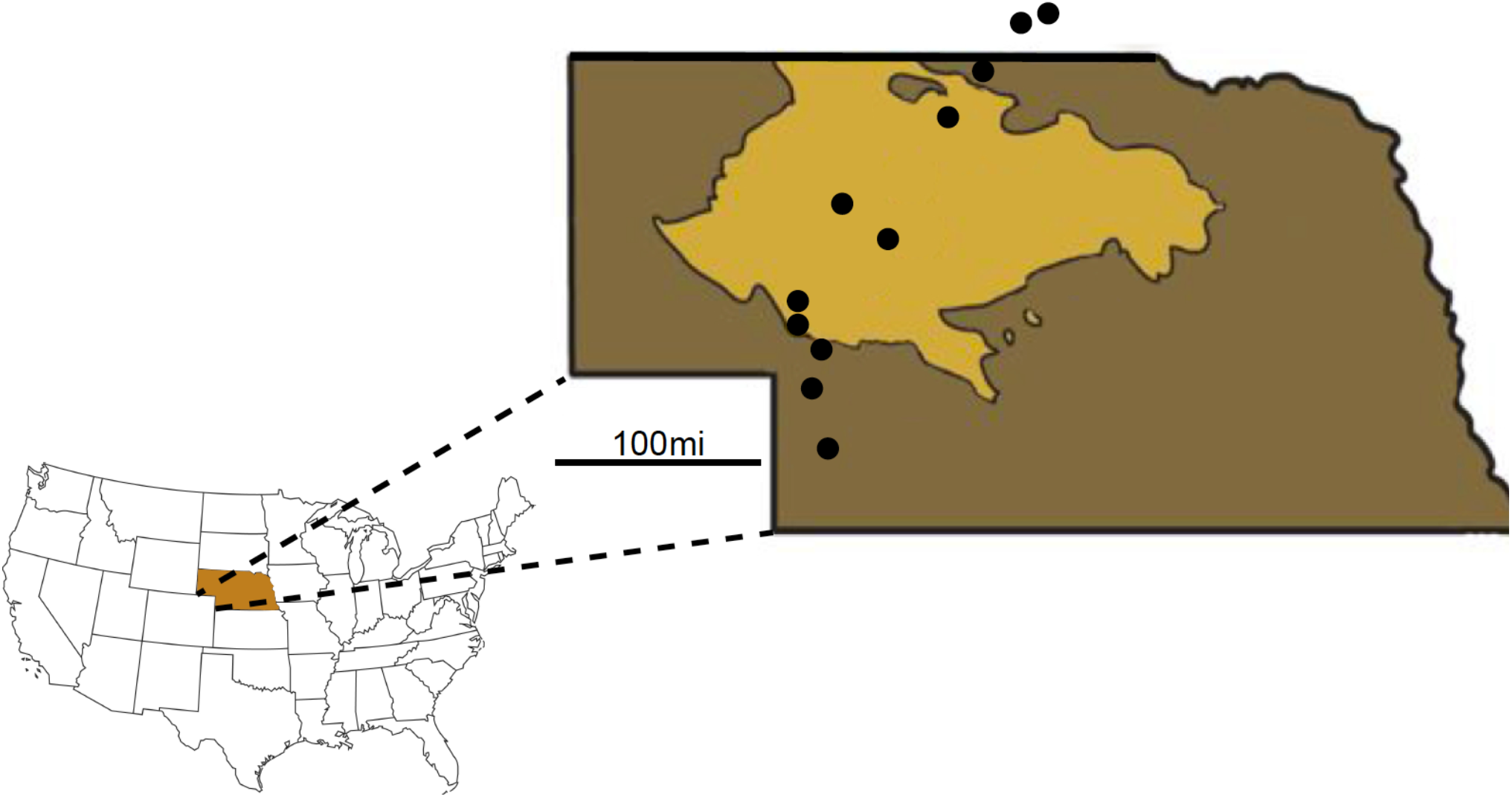
Sampling locations on (light brown) and off (dark brown, and out of state) of the Nebraska Sand Hills. Sampling spanned a 330 km transect beginning ~120 km south of the Sand Hills and ending ~120 km north of the Sand Hills.

**Figure 2.**
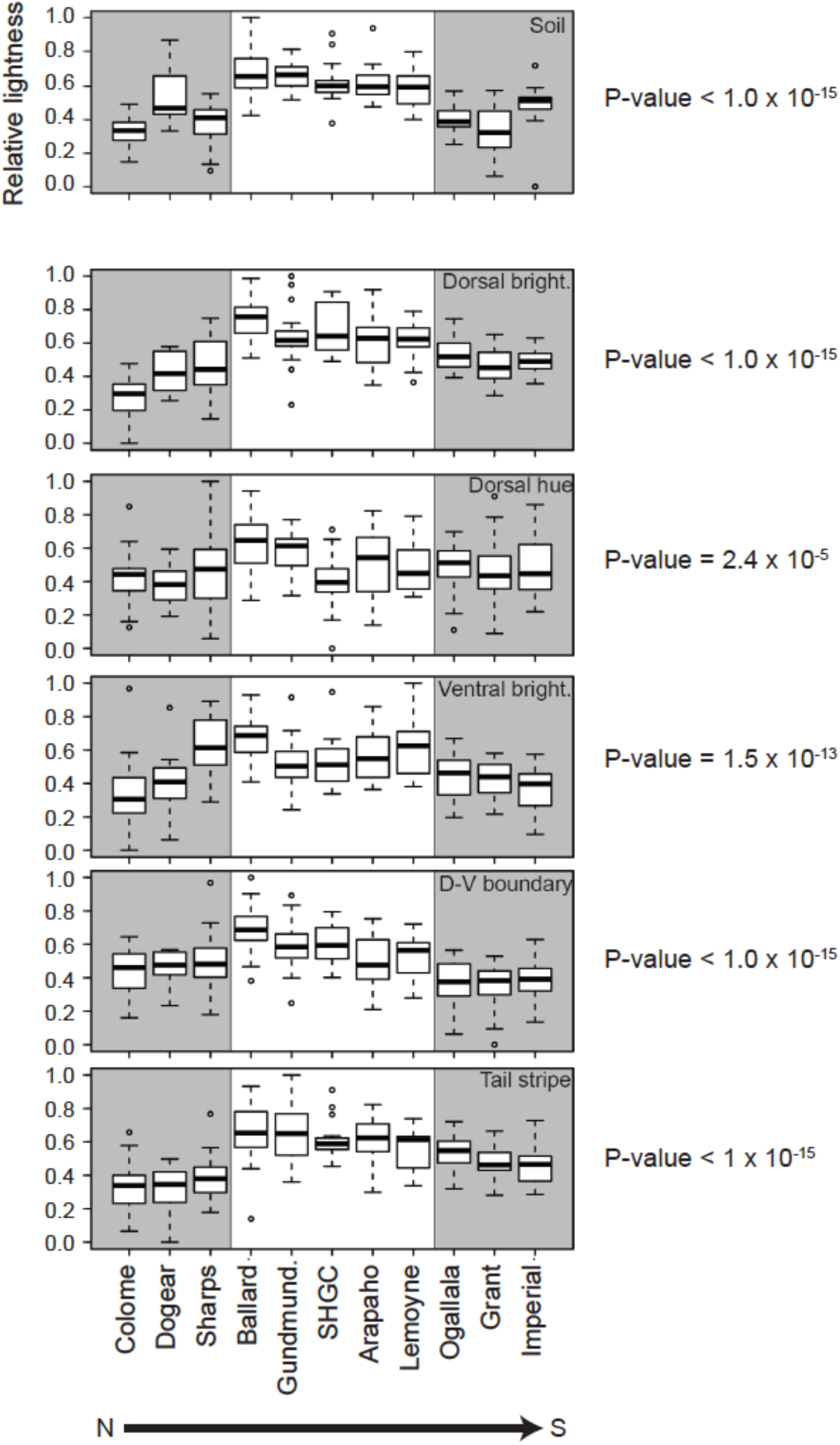
Soil color and mouse color traits are lighter on the Nebraska Sand Hills. Box plots for each sampling site were produced using scaled (range: 0-1), normal-quantile transformed values for: soil brightness (B2), dorsal brightness (PC1), dorsal hue (PC2), ventral brightness (PC3), dorsal-ventral (D-V) boundary, and tail stripe. Soil is shown on the separated top panel, and the five pigment traits are given below. In all plots, higher values correspond to lighter/brighter color (y-axis). Gray shading indicates off the Sand Hills sites. Soil and five mouse color traits were significantly lighter on the Sand Hills (dorsal brightness: *F_1_* = 218. 6, *P* < 1 x 10^−15^; dorsal hue: *F_1_* = 18.4, *P* = 2.4 x 10^−5^; ventral brightness: *F_1_* = 61.0, *P* = 1.5 x 10^−13^; dorsal-ventral boundary: *F_1_* = 104.6, *P* < 1 x 10^−15^; tail stripe: *F_1_* = 170.2, *P* < 1 x 10^−15^), with the corresponding P-values given to the right of each panel. A sixth color trait [ventral hue (PC4); (*F_1_* = 1.1, *P* = 0.30)] and four non-color traits (total body length (*F_1_* = 0.11, *P* = 0.74), tail length (*F_1_* = 0.18, *P* = 0.67), hind foot length (*F_1_* = 3.0, *P* = 0.084), and ear length (*F_1_* = 0.48, *P* = 0.49)) did not differ significantly between habitats (see Supplementary Figure 11, Supplementary Table 11).

We also examined habitat heterogeneity and phenotypic differences among sampling sites within habitat types. Both soil brightness and mouse color differed significantly among sites within habitats (soil: *F_9_* = 8.0, *P* = 1.9 x 10^−10^; dorsal brightness: *F_9_* = 10.5, *P* = 8.1 x 10^−14^; dorsal hue: *F_9_* = 3.4, *P* = 6.7 x 10^−4^; ventral brightness: *F_9_* = 10.3, *P* = 1.4 x 10^−13^; dorsal-ventral boundary: *F_9_* = 6.6, *P* = 1.5 x 10” ^8^; tail stripe: *F_9_* = 6.9, *P* = 7.5 x 10^−9^). Additionally, site-specific means for three of the color traits were significantly correlated with local soil brightness, including: dorsal brightness (*t* = 4.4, *P* = 0.0017), dorsal-ventral boundary (t = 3.9, *P* = 0.0034), and tail stripe (*t* = 3.1, *P* = 0.013). In contrast, site-specific means for dorsal hue and ventral brightness did not correlate with local soil color (dorsal hue: *t* = 2.385, *P* = 0.097; ventral brightness: *t* = 0.32; *P* = 0.77). These results suggest that the agents of selection shaping correlations between local soil color and mouse color vary among the traits.

### The genetic architecture of light color in Sand Hills mice

Genetic architecture parameter estimates derived from our association mapping analyses suggest that variants in the *Agouti* locus explain a considerable amount of the observed color variation in on versus off Sand Hills populations (Table 1). Indeed, *Agouti* SNPs explain 69% of the total phenotypic variance in dorsal brightness and tail stripe. This analysis also provides estimates of the potential number of causal SNPs as well as the proportion of genetic variance that is attributable to major-effect SNPs. These estimates suggest that dozens of *Agouti* variants may be contributing to variation in color traits, but the credible intervals for SNP number are wide (Table 1). One explanation for such wide intervals is that it is difficult to disentangle the effects of closely linked SNPs. Nevertheless, because the lower bound of the credible intervals for all but one color trait exceeds one, our data indicate that there are multiple *Agouti* mutations with non-negligible effects on color phenotypes. Estimates of PGE (percent of genetic variance attributable to major effect mutations) indicate that, whatever their number, these major effect mutations explain a considerable percentage of total genetic variance (*e.g.*, 83% for dorsal brightness and 87% for dorsal-ventral boundary).

**Table 1.**
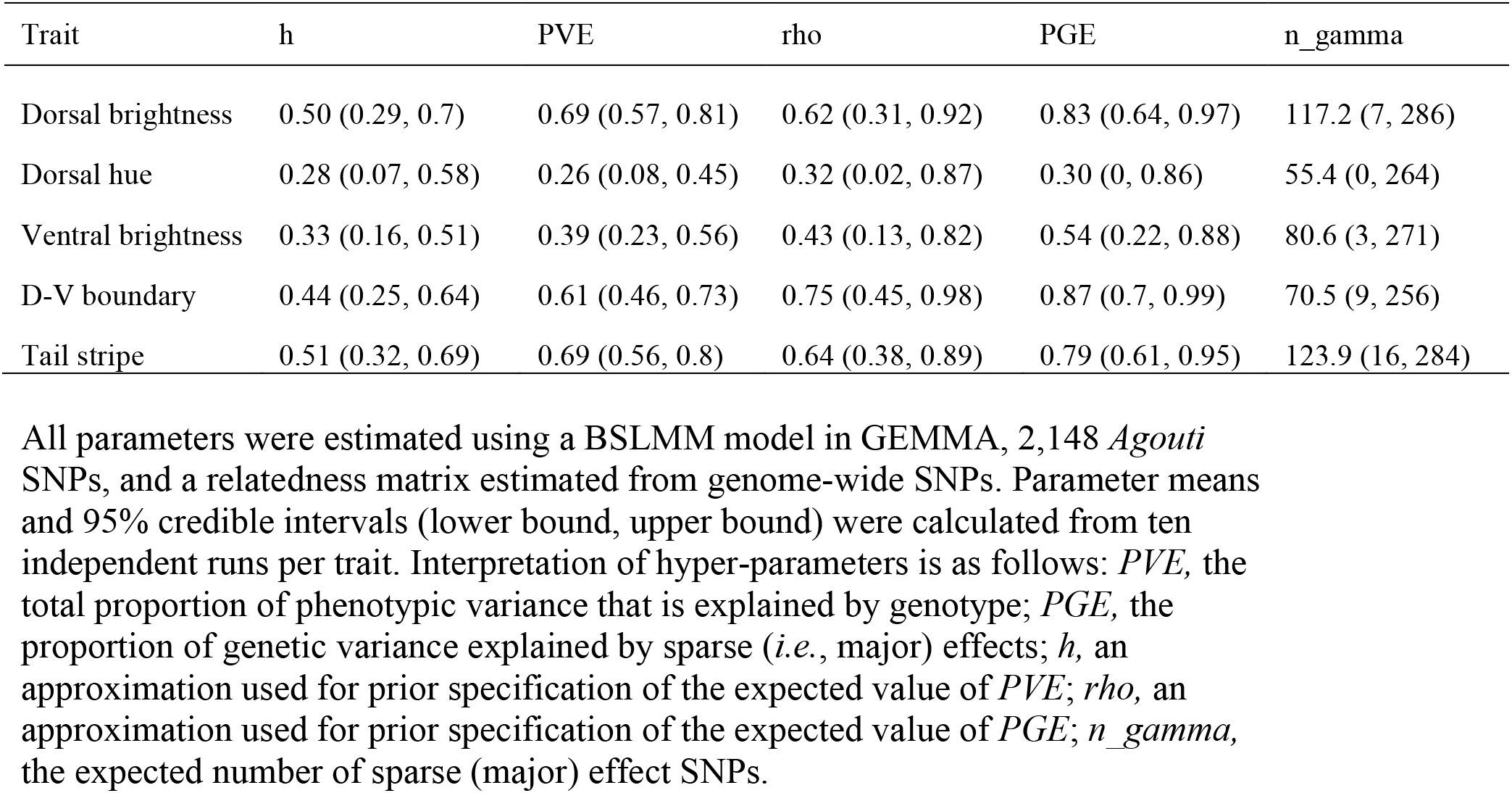
Hyper-parameters describing the genetic architecture of five color traits.

Although our genetic architecture parameter estimates indicate that large-effect *Agouti* variants contribute to variation in each of the five color traits, the number and location of these SNPs vary considerably (Figure 3). To interpret these results, some background on the structure and function of *Agouti* is needed. Work in *Mus musculus* has identified two different *Agouti* isoforms that are under the control of different promoters. The ventral isoform, containing noncoding exons 1A/1A’, is expressed in the ventral dermis during embryogenesis and is required for determining the boundary between the dark dorsum and light ventrum (Bultman *et al*. 1994; Vrieling *et al*. 1994). The hair-cycle isoform, containing noncoding exons 1B/1C, is expressed in both the dorsal and ventral dermis during hair growth, and is required for forming light, pheomelanin bands on individual hairs (Bultman *et al*. 1994; Vrieling *et al*. 1994). In *Peromyscus*, these same isoforms are present as well as additional, novel isoforms (Mallarino *et al*. 2017). Isoform-specific changes in *Agouti* expression are associated with changes in the dorsal-ventral boundary (ventral isoform; Manceau *et al*. 2011) and the width of light bands on individual hairs (hair-cycle isoform; Linnen *et al*. 2009). Both isoforms share the same three coding exons (exons 2, 3, and 4); thus, protein-coding changes could simultaneously alter pigmentation patterning and hair banding (Linnen *et al*. 2013).

**Figure 3.**
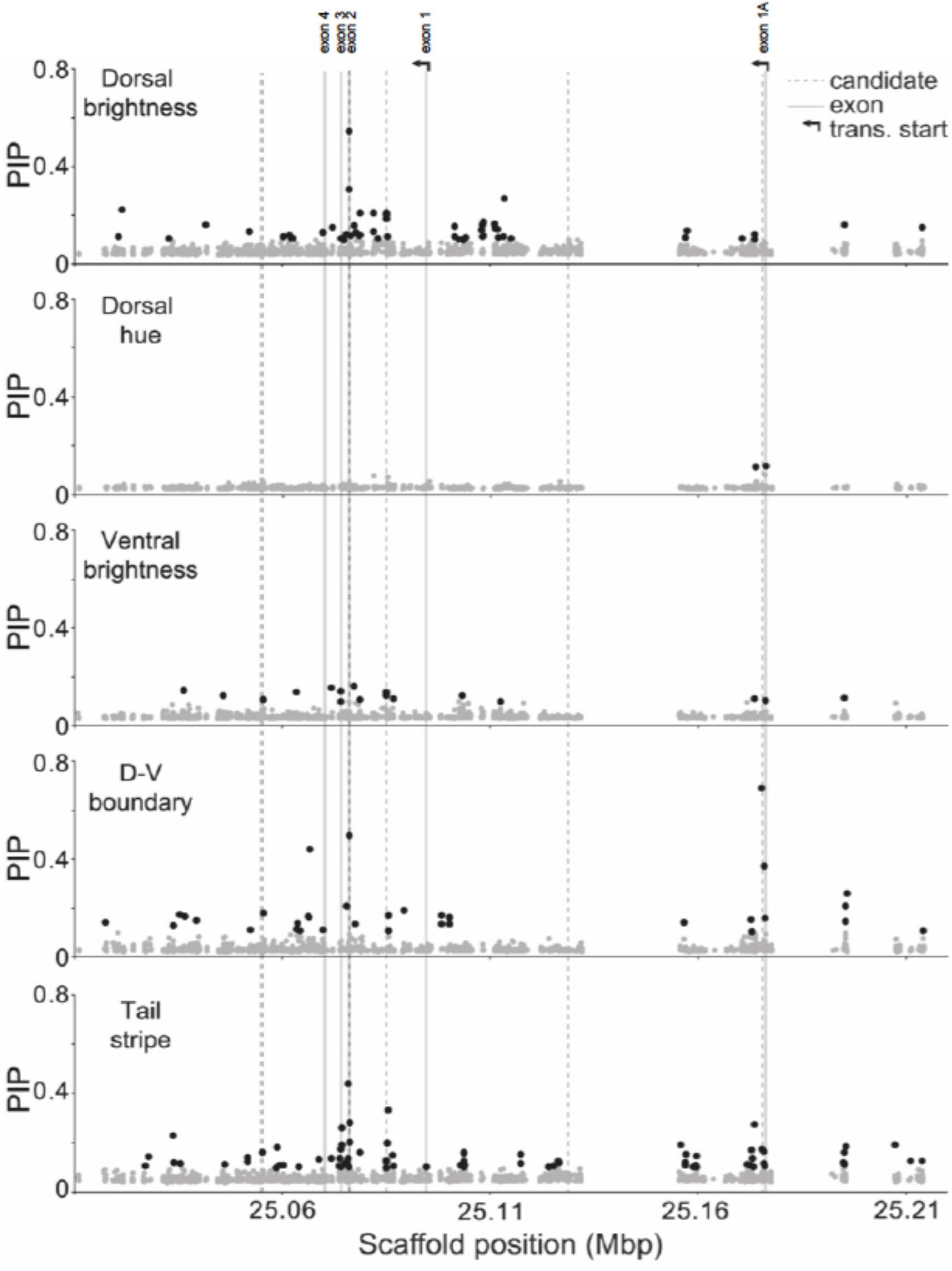
Association mapping results for *Agouti* SNPs and five color traits. Each point represents one of 2,148 *Agouti* SNPs included in the association mapping analysis, with position (on scaffold 16, see “Methods”) indicated on the x-axis and posterior inclusion probabilities (PIP) estimated using a BSLMM in GEMMA on the y-axis. PIP values, which approximate the strength of the association between genotype and phenotype, were averaged across 10 independent GEMMA runs. Black dots indicate SNPs with PIP > 0.1, dashed lines give the location of six candidate SNPs identified in a previous association mapping analysis of a single population (associated with tail stripe, D-V boundary, as well as dorsal brightness and hue; Linnen *et al*. 2013), gray lines give the location of *Agouti* exons, and arrows indicate two alternative transcription start sites.

Across the *Agouti* locus, we detected 160 SNPs that were strongly associated (posterior inclusion probability [PIP] in the polygenic BSLMM > 0.1) with at least one color trait (see Methods). By trait, the number of SNPs exceeding our PIP threshold was: 53 (dorsal brightness), 2 (dorsal hue), 16 (ventral brightness), 34 (dorsal-ventral boundary), and 81 (tail stripe). Additionally, patterns of genotype-phenotype association were distinct for each trait but consistent with *Agouti* isoform function (Figure 3). For example, we observed the strongest associations for dorsal brightness around the three coding exons (exons 2, 3, and 4, located between positions 25.06Mb and 25.11Mb on the scaffold containing the *Agouti* locus) and upstream of the transcription start site for the *Agouti* hair-cycle isoform (Figure 3). By contrast, for dorsal-ventral boundary, the SNP with the highest PIP was located very near the transcription start site for the *Agouti* ventral isoform (Figure 3). Notably, a previously identified serine deletion in exon 2 (Linnen *et al*. 2009; Linnen *et al*. 2013) exceeds our threshold for three of the color traits (dorsal brightness, dorsal-ventral boundary, and tail stripe) and was elevated above background association levels for the remaining two traits (dorsal hue and ventral brightness); and multiple candidates from those analyses are also recovered here (Supplementary Table 2). Overall, our association mapping results thus reveal that there are many sites in *Agouti* that are associated with pigment variation.

We also estimated genetic architecture parameters and identified candidate pigmentation SNPs in the full dataset (*Agouti* SNPs plus all SNPs outside of *Agouti* that had no missing data). For all five traits, including SNPs outside of *Agouti* led to a modest increase in PVE, PGE, and SNP-number estimates, but credible intervals overlapped with those of the *Agouti-only* dataset for all parameter estimates (Table 1 vs. Supplementary Table 3). Additionally, although the highest-PIP SNPs were found in *Agouti*, we identified several *non-Agouti* candidate SNPs as well (Supplementary Table 4). Together, these results suggest that while *Agouti* explains a considerable amount of color variation in mice living on and around the Nebraska Sand Hills, a complete accounting of *non-Agouti* regions associated with color will require a whole-genome resequencing approach.

### The demographic history of the Sand Hills population

Inferring the demographic history of this region is of interest in and of itself, but also is a requisite step for downstream selection inference. Based on their genetic diversity, individuals from different sampling locations grouped according to their location and habitat, with a clear separation between individuals from on and off of the Sand Hills (Figure 4). The observed pattern of isolation by distance was further supported by a significant correlation between pairwise genetic differentiation (*F_ST_*/(1-*F_ST_*)) and pairwise geographic distance among sampling sites (P = 9.9 x 10^−4^). The pairwise *F_ST_* values between sampling locations were low, ranging from 0.008 to 0.065, indicating limited genetic differentiation. Consistent with both patterns of isolation by distance and reduced differentiation among populations, individual ancestry proportions (as inferred by sNMF and TESS) were best explained by three population clusters. Additionally, the ancestry proportions obtained with three clusters (Figure 4) returned low cross-entropy values (Supplementary Figure 1) particularly when accounting for geographic sampling location, again suggesting three distinct groups: (i) north of the Sand Hills, (ii) on the Sand Hills, and (iii) south of the Sand Hills. The population tree inferred with TreeMix also indicated low levels of genetic drift among populations, as well as primary differentiation between north and south populations, with the Sand Hills population occupying an intermediate position. This pattern is consistent with increased differentiation moving along the transect (Figure 4), which is likely owing both to simple isolation by distance as well as the more complex settlement history inferred below.

**Figure 4.**
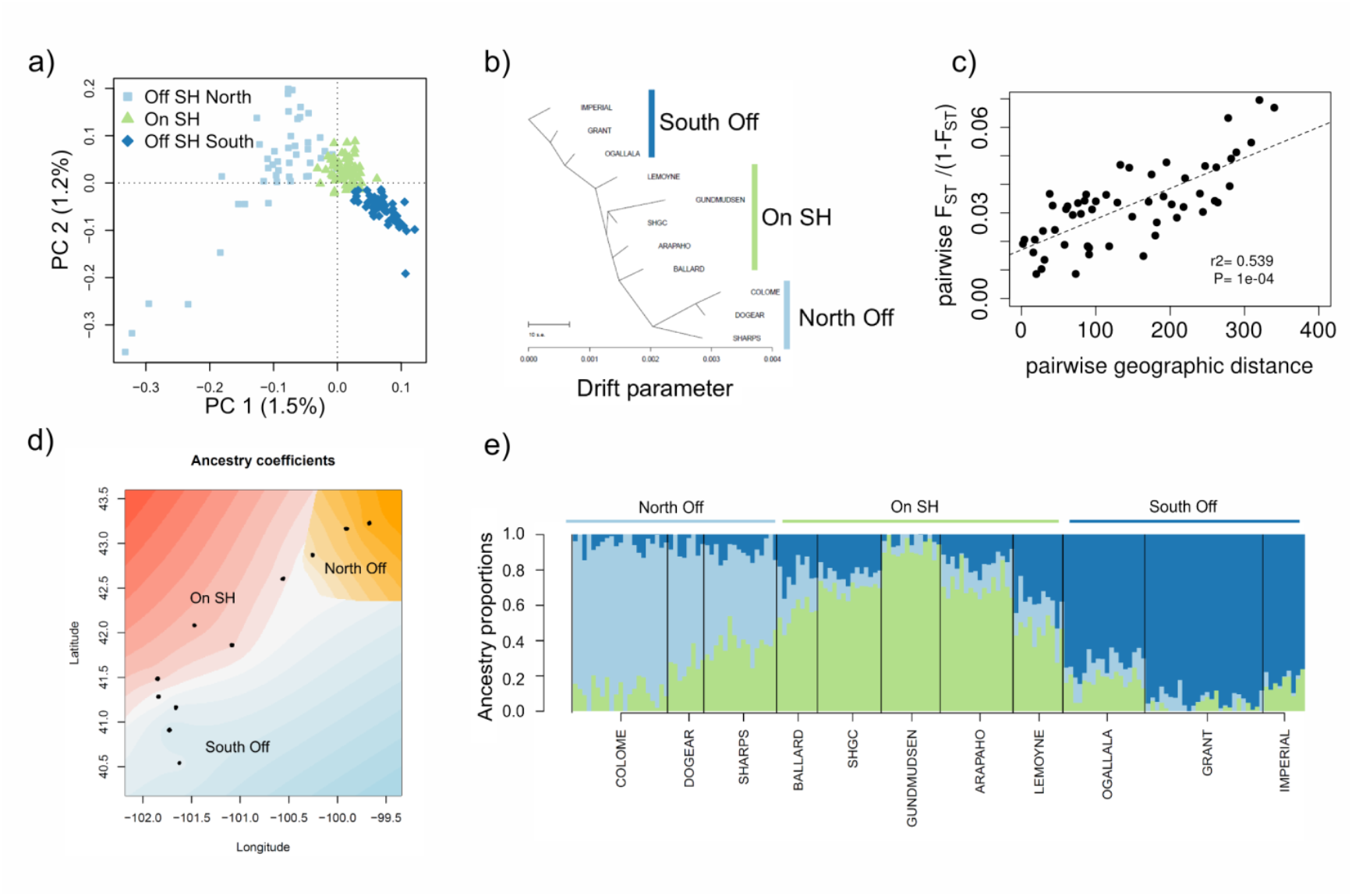
Genetic structure of populations on and off the Sand Hills. **(a)** Principal Component Analysis (PCA) shows that individuals cluster according to their geographic sampling location, with a clear separation of individuals sampled on the Sand Hills (SH; green) from off the Sand Hills (blue). **(b)** TreeMix results for the tree that best fit the covariance in allele frequencies across samples. **(c)** An isolation-by-distance model is supported by a significant correlation between pairwise genetic differentiation (*F_ST_*/(1-*F_ST_*)) and pairwise geographic distance among sampling sites. Significance of the Mantel test was assessed through 10,000 permutations. **(d)** Spatial interpolation of the ancestry proportion inferred, accounting for geographic sampling location, using *TESS3* for *K=3* groups. Individuals are represented as dots, and the ancestry of the three groups is represented by a gradient of three colors. **(e)** Individual admixture proportions (inferred using *TESS3*) are consistent with isolation by distance and reduced differentiation among populations. The number of clusters (*K*) best explaining the data was assessed as the *K* value reaching the lowest minimal crossentropy at *K*=3. All analyses were based on a subset of 161 individuals showing a mean depth of coverage larger than 8X, with all genotypes possessing a minimum coverage of 4X, sampling one SNP for each 1.5kb block, resulting in a dataset with 5,412 SNPs (Supplementary Table 8).

Given the observed population structure, we next investigated models of colonization history based on three populations: those inhabiting the Sand Hills and those inhabiting the neighboring regions to the north and to the south. This resulted in two different three-population demographic models, corresponding to different population tree topologies, which explicitly allow for bottlenecks associated with potential founder events, and for gene flow among populations (Supplementary Figure 2). Both models appeared equally likely (Supplementary Table 5), and parameter estimates were similar and consistent across models, pointing to a recent divergence time among populations, evidence of a bottleneck associated with the colonization of the Sand Hills, and high rates of migration among all populations (Supplementary Table 6). Note that the tested models did not specifically impose a bottleneck for the colonization of the Sand Hills, as population sizes could have remained high at the onset of the colonization of the Sand Hills.

To better distinguish between the models, we compared the likelihoods of the two models for bootstrap datasets containing a single SNP per 1.5kb block, counting the number of bootstrap replicates with a relative likelihood (based on AIC values) larger than 0.95 for each model. Using this approach, we identified a model with a topology in which the population on the Sand Hills shares a more recent common ancestor with the population off the Sand Hills to the south (supported in 81/100 bootstrap replicates, Supplementary Figure 3). This topology and the parameter estimates (Supplementary Table 6) are consistent with a recent colonization of the Sand Hills from the south, namely: (i) a recent split within the last ~4,000 years (95% CI 3,400 to 7,900); (ii) a stronger bottleneck associated with the colonization of the Sand Hills compared to the older split between northern and southern populations; and (iii) higher levels of gene flow from south to north, including higher migration rates from the southern population into the Sand Hills (*2Nm* 95% CI: 12.5-24.0). Furthermore, this model fits well the marginal site frequency spectrum (SFS) for each sample as well as the 2D marginal SFS for each pair of populations (Supplementary Figure 4). Thus, this neutral demographic model nicely explains observed patterns of variation outside of the *Agouti* region, and suggests that, at least the most recent colonization represented by the currently sampled individuals, occurred considerably after the Sand Hills began to initially form (*i.e.*, roughly 8000 years ago).

### The selective history of the Sand Hills population

Despite the high levels of gene flow inferred in our model, which results in low levels of differentiation among populations genome-wide (Supplementary Figure 5a), we observed high levels of differentiation among populations at the *Agouti* locus – variation that is further correlated with several phenotypic traits (Figure 5; Supplementary Figures 5,6). We observed the highest level of differentiation between mice sampled on either side of the northern limit of the Sand Hills, with a 100kb region within *Agouti* exhibiting a continuous run of elevated genetic differentiation (Supplementary Figure 6). This pattern is consistent with the expectations of positive selection under a local adaptation regime. Within this 100kb region, a sub-region of 30kb, located in intron 1 (*i.e.*, between exon 1A and exon 1), displayed maximal differentiation (Supplementary Figure 6), making it difficult to precisely identify the target of selection. In contrast to the wide-range differentiation observed at *Agouti* on the northern side of the cline, a single narrow *F_ST_*-peak of roughly 5kb located in intron 1 was observed on the southern limit of the Sand Hills (Figure 5; Supplementary Figure 6a). Genome-wide comparison confirmed the overall genetic similarity between light-colored Sand Hills mice and the dark-colored population to the south, with the exception of a small number of variants putatively driving adaptive phenotypic divergence at *Agouti* (Figure 5; Supplementary Figure 6a).

**Figure 5.**
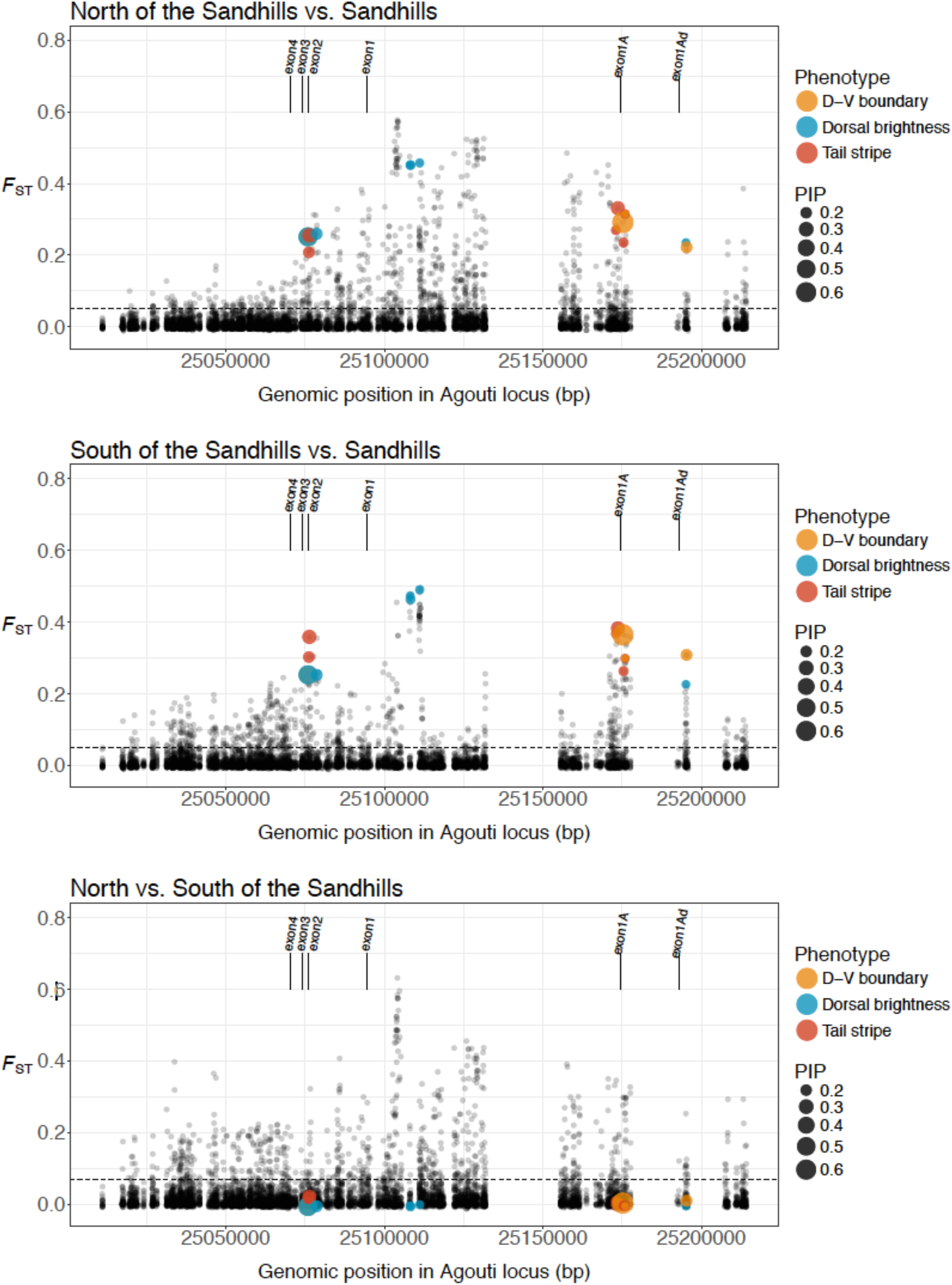
Genetic differentiation across the ***Agouti*** locus. Genetic differentiation observed between populations on the Sand Hills and populations north of the Sand Hills (top panel), populations on the Sand Hills and populations south of the Sand Hills (middle panel), and populations north and south of the Sand Hills. Dotted lines indicate the genome-wide average *F_ST_* between these comparisons. As shown, differentiation is generally greatly elevated across the region, with a number of highly differentiated SNPs particularly when comparing on vs. off Sand Hills populations, which correspond to SNPs associated to different aspects of the cryptic phenotype (PIP scores are indicated by the size of the dots as shown in the legend, and significant SNPs are labeled with respect to the associated phenotype). Populations to the north and south of the Sand Hills are also highly differentiated in this region, likely owing to differing levels of gene flow with the Sand Hills populations, yet the SNPs underlying the cryptic phenotype are not differentiated between these dark populations.

To test whether this pattern of differentiation could be the result of non-selective forces (see Crisci *et al*. 2012; Jensen *et al*. 2016), we calculated the HapFLK statistic, providing a single measure of genetic differentiation for the three geographic localities while controlling for neutral differentiation. Before calculating genetic differentiation at the target locus (*i.e., Agouti*), the HapFLK method computes a neutral distance matrix from the background data (*i.e.*, the background regions randomly distributed across the genome) that summarizes the genetic similarity between populations with regard to the extent of genetic drift since divergence from their common ancestral population (Supplementary Figure 7). Consistent with the inferred demographic history of the populations, genome-wide background levels of variation suggest that individuals captured south of the Sand Hills are more similar to those inhabiting the Sand Hills, when compared with populations to the north. The HapFLK profile confirmed the significant genetic differentiation at the *Agouti* locus compared to neutral expectations (Figure 6; Supplementary Figure 10). Interestingly, southern populations share a significant amount of the haplotypic structure observed in populations on the Sand Hills, with the exception of the candidate region itself. This is in keeping with high rates of on-going gene flow occurring between on and off Sand Hills populations, whereas selection maintains the ancestral *Agouti* alleles in the populations off the Sand Hills, and the derived alleles conferring crypsis on the Sand Hills.

**Figure 6.**
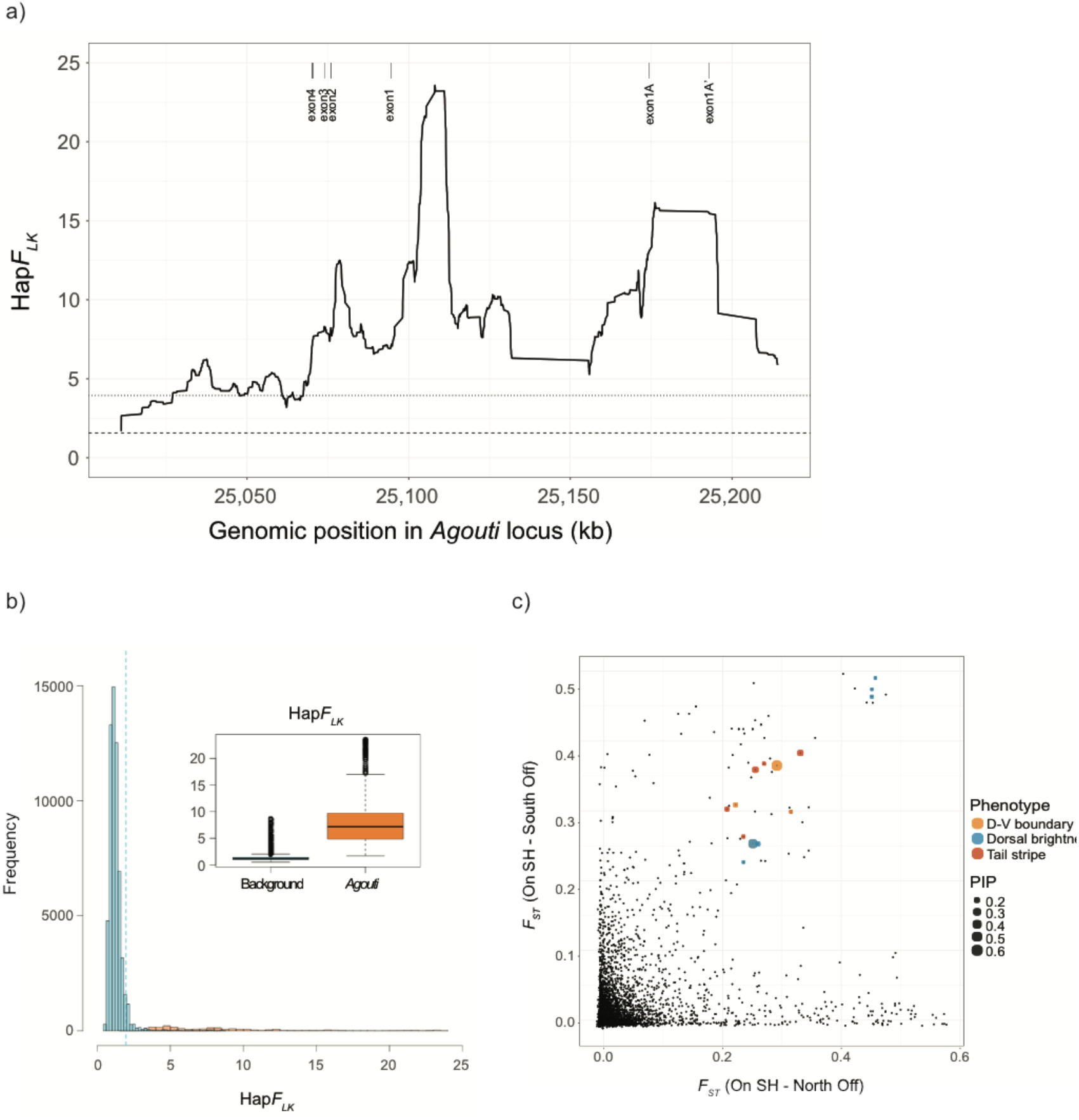
Selection analyses. **(a)** HapFLK profile for the *Agouti* scaffold observed between populations on and off the Sand Hills. Dotted and dashed horizontal lines represent the significance thresholds for the HapFLK statistic using neutral simulations and background data, respectively. **(b)** Distributions of HapFLK values for background (blue) and *Agouti* (orange). The background values are calculated from the genome-wide random background SNPs. Boxplots (insert) are an alternative representation of the two distributions depicted in the main graph. The dashed line indicates the significance threshold using parametric bootstrapping of the best demographic model. **(c)** Genetic differentiation per SNP in *Agouti*. The x-axis represents values obtained from comparing Sand Hills populations to those off of the Sand Hills (north), while the y-axis compares Sand Hills populations to those off of the Sand Hills (south). SNPs associated with phenotypic traits are indicated with their PIP value. Only candidate SNPs with a *F_ST_* value larger than 0.2 in both populations and PIP values larger than 0.15 are included.

Owing to low levels of linkage disequilibrium characterizing the *Agouti* region, adaptive variants could be mapped on a fine genetic scale. Specifically, there are three regions of increased differentiation (Figure 6): (i) a narrow 3kb peak in the HapFLK profile (located in intron 1; Supplementary Table 7) co-localizes with a region of high linkage disequilibrium (Supplementary Figure 8), suggesting a recent selective event; (ii) a second highest peak of differentiation, located between exon 1A and the duplicated reversed copy, exon1A’, is the only significant region detected by the CLR test (Supplementary Figure 9) and contains strong haplotype structure (Figure 6), again suggesting the recent action of positive selection; and (iii) a third highest peak of differentiation surrounds the putatively beneficial serine deletion in exon 2 previously described by Linnen *et al*. (2009), showing strong differentiation between on and off Sand Hills populations in our dataset. Although several lines of evidence support the role of the serine deletion in adaptation to the Sand Hills, the linkage disequilibrium around this variant is low, suggesting that the age of the corresponding selective event is likely the oldest amongst the three candidate regions, with subsequent mutation and recombination events reducing the selective sweep pattern. Hence, as one may anticipate, this greatly expanded clinal data set served to identify younger and more geographically localized candidate regions from across the Sand Hills compared to previous studies, while still generally supporting the model proposed by Linnen *et al*. (2013) of multiple independently selected mutations underlying different aspects of the cryptic phenotype.

### Predicted migration thresholds and the conditions of allele maintenance

Although a number of strong assumptions must be made, it is possible to estimate the migration threshold below which a locally beneficial mutation may be maintained in a population. Given our high rates of inferred gene flow, it is important to consider whether observations are consistent with theoretical expectations necessary for the maintenance of alleles. Following Yeaman & Whitlock (2011) and Yeaman & Otto (2011), this migration threshold is defined in terms of both rates of gene flow as well as fitness effects of locally beneficial alleles in the matching and in the alternate environment. Given the parameter values inferred here, along with estimated population sizes, the threshold above which the most strongly beneficial mutations may be maintained in the population is estimated at *m* = 0.8 – a large value indeed, given the inferred strength of selection (see “Methods” for details). For the more weakly beneficial mutations identified, this value is estimated at *m* = 0.07, again well above empirically estimated rates. Thus, our empirical observations are fully consistent with expectation in that gene flow is strong enough to prevent locally beneficial alleles from fixing on the Sand Hills, but not so strong so as to swamp out the derived allele despite the high input of ancestral variation. As such, parameter estimates fall in a range consistent with the long-term maintenance of polymorphic alleles.

## CONCLUSIONS

The cryptically colored mice of the Nebraska Sand Hills represent one of the few mammalian systems in which aspects of genotype, phenotype, and fitness have been measured and connected. Yet, the population genetics of the Sand Hills region has remained poorly understood. By sequencing hundreds of individuals across a cline spanning over 300 km, fundamental aspects of the evolutionary history of this population could be addressed for the first time. Utilizing genome-wide putatively neutral regions, the inferred demographic history suggests a relatively recent colonization of the Sand Hills from the south well after the last glacial period. Further, high rates of gene flow are inferred between light and dark populations - resulting in genome-wide low levels of genetic differentiation as well as low levels of phenotypic differentiation of four traits unrelated to coloration across the cline. However, the *Agouti* region differs markedly in this regard, with high levels of differentiation observed between on and off Sand Hills populations, strong haplotype structure, and high levels of linkage disequilibrium spanning putatively beneficial mutations among cryptically colored individuals. In addition, we found these putatively beneficial mutations to be strongly associated with the phenotypic traits underlying crypsis, and these phenotypic traits were found to be in strong association with variance in soil color across the cline.

Together, these results suggest a model in which selection is acting to maintain the alleles underlying the locally adaptive phenotype on light/dark soil, despite substantial gene flow, which not only prevents the populations from strongly differentiating from one another, but also prevents the cryptic genotypes from reaching local fixation. Furthermore, returning to the theoretical predictions outlined in the Introduction, the mutations underlying the cryptic phenotype are found to be few in number, of large effect size, and in close genomic proximity. As described by Yeaman & Whitlock (2011) in models of local adaptation with high migration, the establishment of a large effect beneficial allele may indeed facilitate the accumulation of other locally beneficial alleles in the same genomic region owing to the local reduction in effective migration rate (and see Aeschbacher & Burger 2014). Though speculative, such a model may indeed explain the accumulating observations of selection for crypsis generally targeting either the *Agouti* locus or the *Mc1r* locus in a given population, rather than both (i.e., a single region is targeted by selection, rather than two unlinked regions).

Our results are in keeping with this model, with the exon 2 serine deletion explaining a large proportion of variance in multiple traits underlying the cryptic phenotype, and with population-genetic patterns suggesting comparatively old selection acting on this variant. In addition, owing to the large-scale geographic sampling, multiple newly identified, genomically clustered and smaller effect alleles modulating individual traits are also identified, which are characterized by strong patterns of selective sweeps, indicative of more recent selection. This system thus provides an in-depth picture of the dynamic interplay of these population-level processes underlying this adaptive phenotype, and highlights a history characterized by remarkably strong local selective pressures as well as continuous high rates of gene flow with the ancestral founding populations.

## MATERIAL AND METHODS

### Population Sampling

#### Collection

We collected *Peromyscus maniculatus luteus* individuals (*N* = 266) and corresponding soil samples (*N* = 271) from eleven sites spanning a 330 km transect starting ~120 km north of the Sand Hills and ending ~120 km south of the Sand Hills (Figure 1; Supplementary Table 1). In total, five sampling locations were on the Sand Hills and six were located off of the Sand Hills (three in the north and three in the south). We collected mice using Sherman live traps and prepared flat skins according to standard museum protocols; these specimens are accessioned in the Harvard University Museum of Comparative Zoology’s Mammal Department. From each sample, we preserved liver tissue in 100% EtOH for subsequent DNA extraction. Collections were made under the South Dakota Department of Game, Fish, and Parks scientific collector’s permit 54 (2008) and the Nebraska Game and Parks Commission Scientific and Educational Permit (2008) 579, sub-permit 697–700. This work was approved by Harvard University’s Institutional Animal Care and Use Committee.

#### Mouse color measurements

For each mouse, we measured standard morphological traits, including: total length, tail length, hind foot length, and ear length. We also characterized color and color pattern using methods modified from Linnen *et al*. (2013). In brief, to quantify color, we used a USB2000 spectrophotometer (Ocean Optics) to take reflectance measurements at three sites across the body (three replicated measurements in the dorsal stripe, flank, and ventrum). We processed these raw reflectance data using the CLR: Colour Analysis Programs v.1.05 (Montgomerie 2008). Specifically, we trimmed and binned the data to 300-700nm and 1nm bins and computed five color summary statistics: B2 (mean brightness), B3 (intensity), S3 (chroma), S6 (contrast amplitude), and H3 (hue). We then averaged the three measurements for each body region, producing a total of 15 color values (*i.e.*, five summary statistics from each of the three body regions). To ensure values were normally distributed, we performed a normal-quantile transformation on each of the 15 color variables. Using these transformed values, we performed a principal component analysis (PCA) in STATA v.13.0 (StataCorp, College Station, TX). Based on eigenvalues and examination of the scree plot, we selected the first four principal components, which together accounted for 84% of the variation in color. To increase interpretability of the loadings, we performed a VARIMAX rotation on the first four PCs (Tabachnick and Fidell 2001; Montgomerie 2006). After rotation, PC1 corresponded to the brightness/contrast (B2, B3, S6) of the dorsum; PC2 to the chroma/hue (S3, H3) of the dorsum; PC3 to the brightness/contrast (B2, B3, S6) of the ventrum; and PC4 to the chroma/hue (S3, H3) of the ventrum. Therefore, we refer to PC1 as “dorsal brightness”, PC2 as “dorsal hue”, PC3 as “ventral brightness”, and PC4 as “ventral hue” throughout.

To quantify color pattern, we took digital images of each mouse flat skin with a Canon EOS Rebel XTI with a 50mm Canon lens (Canon U.S.A., Lake Success, NY). We used the quick selection tool in Adobe Photoshop CS5 (Adobe Systems, San Jose, CA) to select light and dark areas on each mouse. Specifically, we outlined the dorsum (brown portion of the dorsum with legs and tails excluded), body (outlined entire mouse with brown and white areas, legs and tails excluded), tail stripe (outlined dark stripe on tail only), and tail (outlined entire tail). We calculated “dorsal-ventral boundary” as (body – dorsum)/body and “tail stripe” as (tail – tail stripe)/tail. Thus, each measure represents the proportion of a particular region (tail or body) that appeared unpigmented; higher values therefore represent lighter phenotypes.

To determine whether color phenotypes (dorsal brightness, dorsal hue, ventral brightness, ventral hue, dorsal-ventral boundary, and tail stripe) differ between on Sand Hills and off Sand Hills populations, we performed a nested analysis of variance (ANOVA) on each trait, with sampling site nested within habitat. For comparison, we also performed nested ANOVAs on each of the four non-color traits (total length, tail length, hind foot length, and ear length). All analyses were performed on normal-quantile-transformed data. Unless otherwise noted, we performed all transformations and statistical tests in R v3.3.2.

#### Soil color measurements

To characterize soil color, we collected soil samples in the immediate vicinity of each successful *Peromyscus* capture. To measure brightness, we poured each soil sample into a small petri dish and recorded ten measurements using the USB2000 spectrophotometer. As above, we used CLR to trim and bin the data to 300-700nm and 1nm bins as well as to compute mean brightness (B2). We then averaged these ten values to produce a single mean brightness value for each soil sample, and transformed the soil-brightness values using a normal-quantile transformation prior to analysis. To evaluate whether soil color differs consistently between on and off of the Sand Hills sites, we performed an ANOVA, with sampling site nested within habitat. Finally, to test for a correlation between color traits (see above) and local soil color, we regressed the mean brightness value for each color trait on the mean soil brightness for each sampling site – ultimately for comparison with downstream population genetic inference (see Joost *et al*. 2013).

### Library Preparation and Sequencing

#### Library Preparation

To prepare sequencing libraries, we used DNeasy kits (Qiagen, Germantown, MD) to extract DNA from liver samples. We then prepared libraries following the SureSelect Target Enrichment Protocol v.1.0, with some modifications. In brief, 10-20 μg of each DNA sample was sheared to an average size of 200bp using a Covaris ultrasonicator (Covaris Inc., Woburn, MA). Sheared DNA samples were purified with Agencourt AMPure XP beads (Beckman Coulter, Indianapolis, IN) and quantified using a Quant-iT dsDNA Assay Kit (ThermoFisher Scientific, Waltham, MA). We then performed end repair and adenylation, using 50 μl ligations with Quick Ligase (New England Biolabs, Ipswich, MA) and paired-end adapter oligonucleotides manufactured by Integrated DNA Technologies (Coralville, IA). Each sample was assigned one of 48 five-base pair barcodes. We pooled samples into equimolar sets of 9-12 and performed size selection of a 280-bp band (+/− 50bp) on a Pippin Prep with a 2% agarose gel cassette. We performed 12 cycles of PCR with Phusion High-Fidelity DNA Polymerase (NEB), according to manufacturer guidelines. To permit additional multiplexing beyond that permitted by the 48 barcodes, this PCR step also added one of 12 six-base pair indices to each pool of twelve barcoded samples. Following amplification and AMPure purification, we assessed the quality and quantity of each pool (23 total) with an Agilent 2200 TapeStation (Agilent Technologies, Santa Clara, CA) and a Qubit-BR dsDNA Assay Kit (Thermo Fisher Scientific, Waltham, MA).

#### Enrichment and Sequencing

To enrich sample libraries for both the *Agouti* locus as well as more than 1000 randomly distributed genomic regions, and following Linnen *et al*. (2013), we used a MYcroarray MYbaits capture array (MYcroarray, Ann Arbor, MI). This probe set includes the 185kb region containing all known *Agouti* coding exons and regulatory elements (based on a *P. maniculatus rufinus Agouti* BAC clone from Kingsley *et al*. 2009) and >1000 non-repetitive regions averaging 1.5kb in length at random locations across the *P. maniculatus* genome.

After generating 23 indexed pools of barcoded libraries from 249 individual Sand Hills mice and one lab-derived non-agouti control, we enriched for regions of interest following the standard MYbaits v.2 protocol for hybridization, capture, and recovery. We then performed 14 cycles of PCR with Phusion High-Fidelity Polymerase and a final AMPure purification. Final quantity and quality was then assessed using a Qubit-HS dsDNA Assay Kit and Agilent 2200 TapeStation. After enriching our libraries for regions of interest, we combined them into four pools and sequenced across eight lanes of 125bp paired-end reads using an Illumina HiSeq2500 platform (Illumina, San Diego, CA). Of the read pairs, 94% could be confidently assigned to individual barcodes (*i.e.*, we excluded read data where the ID tags were not identical between the two reads of a pair (4%) as well as reads where ID tags had low base qualities (2%)). Read pair counts per individual ranged from 367,759 to 42,096,507 (median 4,861,819).

### Reference Assembly

We used the *Peromyscus maniculatus bairdii* Pman_1.0 reference assembly publicly available from NCBI (RefSeq assembly accession GCF_000500345.1), which consists of 30,921 scaffolds (scaffold N50: 3,760,915) containing 2,630,541,020bp ranging from 201bp to 18,898,765bp in length (median scaffold length: 2,048bp; mean scaffold length: 85,073bp) to identify variation in the background regions (*i.e.*, outside of *Agouti*). Unfortunately, the *Agouti* locus is split over two overlapping scaffolds (*i.e.*, exon 1A/A’ is located on scaffold NW_006502894.1 and exon 2, 3, and 4 are located on scaffold NW_006501396.1) in this assembly, causing issues in the read mapping at this locus. Therefore, reads mapped on either of these two scaffolds were re-aligned to a novel in-house *Peromyscus* reference assembly in order to more reliably identify variation at the *Agouti* locus. The in-house reference assembly consists of 9,080 scaffolds (scaffold N50: 13,859,838) containing 2,512,380,343bp ranging from 1,000bp to 60,475,073bp in length (median scaffold length: 3,275bp; mean scaffold length: 276,694bp). In the two reference assemblies, we annotated and masked seven different classes of repeats (*i.e.*, SINE, LINE, LTR elements, DNA elements, satellites, simple repeats, and low complexity regions) using RepeatMasker v.Open-4.0.5 (Smit *et al*. 2013).

### Sequence Alignment

We removed contamination from the raw read pairs and trimmed low quality read ends using cutadapt v.1.8 (Martin 2011) and TrimGalore! v.0.3.7 (http://www.bioinformatics.babraham.ac.uk/projects/trim_galore). We aligned the preprocessed, paired-end reads to the reference assembly using Stampy v.1.0.22 (Lunter and Goodson 2011). PCR duplicates as well as reads that were not properly paired were then removed using SAMtools v.0.1.19 (Li *et al*. 2009). After cleaning, read pair counts per individual ranged from 220,960 to 20,178,669 (median: 3,062,998). We then conducted a multiple sequence alignment using the Genome Analysis Toolkit (GATK) IndelRealigner v.3.3 (McKenna *et al*. 2010; DePristo *et al*. 2011; Van der Auwera *et al*. 2013) to improve variant calls in low-complexity genomic regions, adjusting Base Alignment Qualities (BAQ) at the same time in order to down weight base qualities in regions with high ambiguity (Li 2011). Next, we merged sample-specific reads across different lanes, thereby removing optical duplicates using SAMtools v.0.1.19. Following GATK’s Best Practice, we performed a second multiple sequence alignment to produce consistent mappings across all lanes of a sample. The resulting dataset contained aligned read data for 249 individuals from 11 different sampling locations.

### Variant Calling and Filtering

We performed an initial variant call using GATK’s HaplotypeCaller v.3.3 (McKenna *et al*. 2010; DePristo *et al*. 2011; Van der Auwera *et al*. 2013), and then we genotyped all samples jointly using GATK’s GenotypeGVCFs v.3.3 tool. Post-genotyping, we filtered initial variants using GATK’s VariantFiltration v.3.3, removing SNPs using the following set of criteria (with acronyms as defined by the GATK package): (i) the variant exhibited a low read mapping quality (MQ<40); (ii) the variant confidence was low (QD<2.0); (iii) the mapping qualities of the reads supporting the reference allele and those supporting the alternate allele were qualitatively different (MQRankSum<-12.5); (iv) there was evidence of a bias in the position of alleles within the reads that support them between the reference and alternate alleles (ReadPosRankSum<-8.0); or (v) there was evidence of a strand bias as estimated by Fisher’s exact test (FS>60.0) or the Symmetric Odds Ratio test (SOR>4.0). We removed indels using the following set of criteria: (i) the variant confidence was low (QD<2.0); (ii) there was evidence of a strand bias as estimated by Fisher’s exact test (FS>200.0); or (iii) there was evidence of bias in the position of alleles within the reads that support them between the reference and alternate alleles (ReadPosRankSum<-20.0). For an in-depth discussion of these issues, see the recent review of Pfeifer (2017).

To minimize genotyping errors, we excluded all variants with a mean genotype quality of less than 20 (corresponding to P[error] = 0.01). We limited SNPs to biallelic sites using VCFtools v.0.1.12b (Danecek *et al*. 2011), with the exception of one previously identified putatively beneficial triallelic variant (Linnen *et al*. 2013). We excluded variants within repetitive regions of the reference assembly from further analyses as potential mis-alignment of reads to these regions might lead to an increased frequency of heterozygous genotype calls. Additionally, we filtered variants on the basis of the Hardy-Weinberg equilibrium (HWE) by computing a p-value for HWE for each variant using VCFtools v.0.1.12b and excluding variants with an excess of heterozygotes (*p* < 0.01), unless otherwise noted. Finally, we removed sites for which all individuals were fixed for the non-reference allele as well as sites with more than 25% missing genotype information.

The resulting call set contained 8,265 variants on scaffold 16 (containing *Agouti)* and 649,300 variants within the random background regions. We polarized variants within *Agouti* using the *P. maniculatus rufinus Agouti* sequence (Kingsley *et al*. 2009) as the outgroup. Genotypes were phased using BEAGLE v.4 (Browning and Browning 2007).

### Accessible Genome

To minimize the number of false positives in our dataset, we subjected variants to several stringent filter criteria. The application of these filters led to an exclusion of a substantial fraction of genomic regions, and thus we generated mask files (using the GATK pipeline described above, but excluding variant-specific filter criteria *i.e.*, QD, FS, SOR, MQRankSum, and ReadPosRankSum) to identify all nucleotides accessible to variant discovery. After filtering, only a small fraction of the genome (*i.e.*, 63,083bp within *Agouti* and 6,224,355bp of the random background regions) remained accessible. These mask files enabled us to obtain an exact number of the monomorphic sites in the reference assembly, which we then used to both estimate the demographic history of focal populations as well as a control to avoid biases when calculating summary statistics.

### Association mapping

To identify SNPs within *Agouti* that contribute to variation in mouse coat color, we used the Bayesian sparse linear mixed model (BSLMM) implemented in the software package GEMMA v 0.94.1 (Zhou *et al*. 2013). In contrast to single-SNP association mapping approaches (*e.g.*, Purcell *et al*. 2007), the BSLMM is a polygenic model that simultaneously assesses the contribution of multiple SNPs to phenotypic variation. Additionally, compared to other polygenic models (*e.g.*, linear mixed models and Bayesian variable selection regression), the BSLMM performs well for a wide range of trait genetic architectures, from a highly polygenic architecture with normally distributed effect sizes to a “sparse” model in which there are a small number of large-effect mutations (Zhou *et al*. 2013). Indeed, one benefit of the BSLMM (and other polygenic models) is that hyper-parameters describing trait genetic architecture (*e.g.*, number of SNPs, relative contribution of large- and small-effect variants) can be estimated from the data. Importantly, the BSLMM also accounts for population structure via inclusion of a kinship matrix as a covariate in the mixed model.

For each of the five color traits that differed significantly between on Sand Hills and off Sand Hills habitats (*i.e.*, dorsal brightness, dorsal hue, ventral brightness, dorsal-ventral boundary, and tail stripe, see “Results and Discussion”), we performed ten independent GEMMA runs, each consisting of 25 million generations, with the first five million generations discarded as burn-in. Because we were specifically interested in the contribution of *Agouti* variants to color variation, we restricted our GEMMA analysis to 2,148 *Agouti* SNPs that had no missing data and a minor allele frequency (MAF) > 0.05. However, to construct a relatedness matrix, we used the genome-wide SNP dataset. We aimed to maximize the number of individuals kept in the analyses, and hence we used less stringent filtering criteria than for the demographic analyses. Prior to construction of this matrix, we removed all individuals with a mean depth (DP) of coverage lower than 2X. Following the removal of low-coverage individuals, we treated genotypes with less than 2X coverage or twice the individual mean DP as missing data and removed SNPs with more than 10% missing data across individuals or with a MAF<0.01. To remove tightly linked SNPs, we sampled one SNP per 1kb block, choosing whichever SNP had the lowest amount of missing data, resulting in a dataset with 12,920 SNPs (Supplementary Table 8). We then computed the relatedness matrix using the R/Bioconductor (release 3.4) package SNPrelate v1.8.0 (Zheng *et al*. 2012). For all traits, we used normal quantile transformed values to ensure normality, and option “-bslmm 1” to fit a standard linear BSLMM to our data.

After runs were complete, we assessed convergence on the desired posterior distribution via examination of trace plots for each parameter and comparison of results across independent runs. We then summarized posterior inclusion probabilities (PIP) for each SNP for each trait by averaging across the ten independent runs. Following (Chaves *et al*. 2016), we used a strict cut-off of PIP>0.1 to identify candidate SNPs for each color trait (cf. Gompert *et al*. 2013; Comeault *et al*. 2014). To summarize genetic architecture parameter estimates for each trait, we calculated the mean and upper and lower bounds of the 95% credible interval for each parameter from the combined posterior distributions derived from the 10 runs.

To identify additional candidate regions contributing to color variation and to compare genetic architecture parameter estimates from *Agouti* to those obtained using the full dataset (*Agouti* SNPs and *non-Agouti* SNPs), we repeated the GEMMA analyses as described above, but with a dataset consisting of 8,616 SNPs that had no missing data and MAF > 0.05.

### Population Structure

We investigated the structure of populations along the Nebraska Sand Hills transect with several complementary approaches. First, we used methods to cluster individuals based on their genetic similarity, including PCA and inference of individual ancestry proportions. Second, on the basis of SNP allele frequencies, we computed pairwise *F_ST_* between sampled sites to infer their relationships. In addition, we inferred the population tree best fitting the covariance of allele frequencies across sites using TreeMix v1.13 (Pickrell & Pritchard 2012). Finally, we tested for isolation by distance (IBD) by comparing the matrix of geographic distances between sites to that of pairwise genetic differentiation (*F_ST_*/(1-*F_ST_*) (Rousset 1997)) using Mantel tests (Sokal 1979; Smouse *et al*. 1986) as well as methods to infer ancestry proportions accounting for geographic location.

For demographic analyses, we based all analyses on the background regions randomly distributed across the genome (*i.e.*, excluding *Agouti*) and applied additional filters to maximize the quality of the data. First, we discarded individuals with a mean depth of coverage (DP) across sites lower than 8X. Second, given that the DP was heterogeneous across individuals, we treated all genotypes with DP<4 or with more than twice the individual mean DP as missing data to avoid genotype call errors due to low coverage or mapping errors. Third, we partitioned each scaffold into contiguous blocks of 1.5kb size, recording the number of SNPs and accessible sites for each block. To minimize regions with spurious SNPs (e.g., due to mis-alignment or repetitive regions), we only kept blocks with more than 150bp of accessible sites and with a median distance among consecutive SNPs larger than 3bp. This resulted in a dataset consisting of 190 individuals with ~2.8Mb distributed across 11,770 blocks, with 284,287 SNPs and a total of 2,814,532 callable sites (corresponding to 2,530,245 monomorphic sites; Supplementary Table 9). Fourth, to minimize missing data, we only kept SNPs with at least 90% of called genotypes, and individuals with at least 75% of data across sites in the dataset. Finally, since many of the applied methods rely on the assumption of independence among SNPs, we generated a dataset by sampling one SNP per block, selecting the SNP with the lowest amount of missing data.

The PCA and the pairwise *F_ST_* (estimated following Weir & Cockerham 1984) analyses were performed in R using the method implemented in the Bioconductor (release 3.4) package SNPrelate v1.8.0 (Zheng *et al*. 2012). We inferred the ancestry proportions of all individuals based on *K* potential components using sNMF (Frichot *et al*. 2014) implemented in the R/Bioconductor release 3.4 package LEA v1.6.0 (Frichot & François 2015) with default settings. We examined *K* values from 1 to 12, and selected the *K* that minimized the cross entropy as suggested by Frichot *et al*. (2014). We performed the PCA, *F_ST_* and sNMF analyses by applying an extra minor allele frequency (MAF) filter > (1/2n), where *n* is the number of individuals, such that singletons were discarded. To test for isolation by distance, we compared the estimated *FST* values to the pairwise geographic distances (measured as a straight line from the most northern sample, *i.e.*, Colome) between sample sites using a Mantel test, with significance assessed by 10,000 permutations of the genetic distance matrix. In addition, fine scale population structure was accounted for using the TESS3 R package (Caye *et al*. 2016). Given the similar cross-entropy values in sNMF for *K* = 1 to 3, 100 independent runs for each *K* value were performed with a tolerance of 10^−6^ and 200 iterations per run.

### Demographic Analyses

To uncover the colonization history of the Nebraska Sand Hills, we inferred the demographic history of populations based on the joint site frequency spectrum (SFS) obtained from the random anonymous genomic regions, using the composite likelihood method implemented in fastsimcoal2 v3.05 (Excoffier *et al*. 2013). In particular, we were interested in testing whether the Sand Hills populations were founded widely from across the ancestral range, or whether there was a single colonization event. We also aimed to quantify the current and past levels of gene flow among populations. We considered models with three populations corresponding to samples on the Sand Hills (*i.e.*, Ballard, Gudmundsen, SHGC, Arapaho, and Lemoyne) as well as off of the Sand Hills in the north (*i.e.*, Colome, Dogear, and Sharps) and in the south (*i.e.*, Ogallala, Grant, and Imperial). Specifically, we considered two alternative three-population demographic models, as those are best supported by the data, to test whether the colonization of the Sand Hills most likely occurred from the north or from the south, in a single event or serial events, thereby simultaneously quantifying the levels of gene flow between populations on and off of the Sand Hills (Supplementary Figure 2). In these models, we assumed that colonization dynamics were associated with founder events (*i.e.*, bottlenecks), and the number and place of which along the population trees were allowed to vary among models. However, parameter values corresponding to a no size change model (*i.e.*, no bottleneck) were included for evaluation. Mutation rates and generation times were taken following Linnen *et al*. (2009, 2013).

We constructed a three-dimensional (3D) SFS by pooling individuals from the three sampling regions (*i.e.*, on the Sand Hills as well as off of the Sand Hills in the north and south). For each of the 11,770 blocks of 1.5kb size, 30 individuals (*i.e.*, ten individuals from each of the three geographic regions) were selected such that all SNPs within a given block exhibited complete genotype information. Specifically, we selected the 30 individuals with the least missing data for each block, only including SNPs with complete genotype information across the chosen individuals, a procedure that maximized the number of SNPs without missing data while keeping the local pattern of linkage disequilibrium within each block. Note that we sampled genotypes of the same individuals within each block, but that at different blocks the sampled individuals might differ. For each block, we computed the number of accessible sites and discarded blocks without any SNPs.

The resulting 3D-SFS contained a total of 140,358 SNPs and 2,674,174 accessible sites (Supplementary Table 8). The number of monomorphic sites was based on the number of accessible sites, assuming that the proportion of SNPs lost with the extra DP filtering steps, not included in the mask file, was identical for polymorphic and monomorphic sites. Given that there was no closely related outgroup sequence available to determine the ancestral allele for SNPs within the random background regions, we analyzed the multidimensional folded site frequency spectrum, generated using Arlequin v.3.5.2.2 (Excoffier & Lischer 2010). For each model, we estimated the parameters that maximized its likelihood by performing 50 optimization cycles (-L 50), approximating the expected SFS based on 350,000 coalescent simulations (-N 350,000), and using as a search range for each parameter the values reported in Supplementary Table 10.

The model best fitting the data was selected using Akaike’s information criterion (AIC; Akaike 1974). Because our dataset likely contains linked sites, the confidence for a given model is likely to be inflated. Thus, to compare models, we generated bootstrap replicates with one SNP sampled from each 1.5kb block, which we assumed to be independent. We estimated the likelihood of each bootstrap replicate based on the expected SFS obtained for each model with the full set of SNPs (following Bagley *et al*. 2017). Furthermore, we obtained confidence intervals for the parameter estimates by a non-parametric block-bootstrap approach. To account for linkage disequilibrium, we generated 100 bootstrap datasets by sampling the 11,770 blocks for each bootstrap dataset with replacement in order to match the original dataset size. For each dataset, we performed two independent runs using the parameters that maximized the likelihood as starting values. We then computed the 95 percentile confidence intervals of the parameters using the R-package boot v.1.3-18.

### Selection Analyses

We used VCFtools v.0.1.12b (Danecek *et al*. 2011) to calculate Weir and Cockerham’s *F_ST_* (Weir & Cockerham 1984) in the *Agouti* region. We pooled the 11 sampling locations into three populations (*i.e.*, one population on the Sand Hills and two populations off of the Sand Hills: one in the north and one in the south, see “Demographic Analyses”) and calculated *F_ST_* in a sliding window (window size 1kb, step size 100bp) as well as on a per-SNP basis within *Agouti* to identify highly differentiated candidate regions for local adaptation.

To control for potential hierarchical population structure as well as past fluctuations in population size, we also measured genetic differentiation using the HapFLK method (Fariello *et al*. 2013). HapFLK calculates a global measure of differentiation for each SNP (FLK) or inferred haplotype (HapFLK) after having rescaled allele/haplotype frequencies using a population kinship matrix. We calculated the kinship matrix using the complete dataset, excluding the scaffolds containing *Agouti*. We launched the HapFLK software using 40 independent runs (--nfit 40) and −K 40, only keeping alleles with a MAF > 0.05. We obtained the neutral distribution of the HapFLK statistic by running the software on 1,000 neutral simulated datasets under our best demographic model.

To map potential complete selective sweeps in the *Agouti* region, we utilized the CLR method (Nielsen *et al*. 2005) as implemented in the software Sweepfinder2 (DeGiorgio *et al*. 2016). For the analysis, we used the *P. maniculatus rufinus Agouti* sequence to identify ancestral and derived allelic states (see “Variant Calling and Filtering”). We ran Sweepfinder2 with the “–su” option, defining grid-points at every variant and using a pre-computed background SFS. We calculated the cutoff value for the CLR statistic using a parametric bootstrap approach (as proposed by Nielsen *et al*. 2005). For this purpose, we re-ran the CLR analysis on 10,000 datasets simulated under our inferred neutral demographic model for the Sand Hills populations in order to reduce false-positive rates (see Crisci *et al*. 2013; Poh *et al*. 2014).

### Calculating Conditions of Allele Maintenance

Following Yeaman & Whitlock (2011), the threshold for allele maintenance is defined as the migration rate that satisfies *δ*=1/(4*N*), which represents the criteria at which allele frequency changes owing to genetic drift are on the same order as frequency changes owing to the interplay between selection and migration. Further, in order to extend this migration threshold to the consideration of a phenotypic trait, they define the fitness (*W*) of phenotype (*Z*) as:

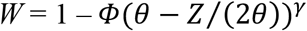

where *Φ* is the strength of stabilizing selection, *θ* is the locally adaptive phenotype which takes a positive value in the derived population and a negative value in the ancestral population, and *γ* specifies a curvature for the function. Further, the effect size of the underlying mutation is given as *α*. Following Yeaman & Otto (2011), *δ* may then be defined as:

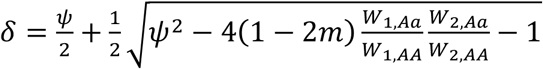

where 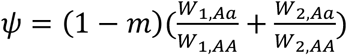, *W_ij_*, is the fitness of allele *j* in environment *i*, *a* is the allele beneficial in the ancestral environment and deleterious in the derived environment, and *A* is the allele that is beneficial in the derived environment. Finally, Yeaman & Whitlock (2011) define the migration threshold for a particular value of *α* as:

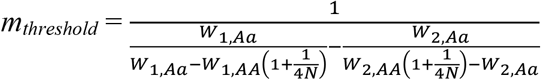

To compare with our empirical observations and inferred values, we take for the sake of example the identified serine deletion in exon 2, compared between the Sand Hills and the northern population. Firstly, we set *θ* = ±1 (*i.e.*, positive on the Sand Hills, negative off of the Sand Hills), and *γ* = 2 (*i.e.*, a convex shape (Yeaman & Whitlock 2011)). Inference pertaining to the strength of selection acting on the cryptic phenotype has been made, most notably from previous predation experiments in which conspicuously colored phenotypes were attacked significantly more often than those that were cryptically colored, with a calculated selection index = 0.545 (Linnen *et al*. 2013). Furthermore, previous crossing experiments have suggested that the serine deletion is a dominant allele (Linnen *et al*. 2009). Finally, the most strongly associated trait has here been calculated near 0.5 (i.e., dorsal brightness). For the corresponding value of *α*, this suggests a migration threshold of *m* = 0.8. Thus, for the serine deletion, it is readily apparent that our inferred migration rates are well below this expected threshold allowing maintenance (where the 95% CI is contained in *m* < 0.0004).

## ACKNOWLEDGEMENTS

We thank J. Larson and K. Turner for laboratory assistance; E. Kay, E. Kingsley, and M. Manceau for field assistance; J. Demboski and the Denver Museum of Nature and Science for logistical support; the University of Nebraska-Lincoln for use of facilities and/or permission to collect mice at Cedar Point Biological Station, Gudmundsen Sandhills Laboratory, and Arapaho Prairie; and J. Chupasko for curation assistance. This work was funded by a Swiss National Science Foundation Sinergia grant to LE, HEH, and JDJ.

---

**Supplementary Table 1.** Sampling locations and sample sizes for mice and soil samples.

| Site name | Location | Latitude | Longitude | Habitat | N<br>(pheno) | N<br>(soil) | N<br>(geno) |
| --- | --- | --- | --- | --- | --- | --- | --- |
| Colome | South Dakota: Tripp Co.,<br>cornfield near Colome, SD | N43°13.485' | W99°40.353' | Off | 30 | 30 | 31 |
| Dogear | South Dakota: Tripp Co.,<br>near Dog Ear Lake | N43°09.766' | W99°54.581' | Off | 13 | 15 | 14 |
| Sharps | Nebraska: Cherry Co.,<br>Sharp's Campground, Sparks,<br>NE | N42°52.709' | W100°15.621' | Off | 33 | 31 | 33 |
| Ballard | Nebraska: Cherry Co.,<br>Ballards Marsh State Wildlife<br>Management Area, Valentine,<br>NE | N42°36.147' | W100°33.733' | SH | 33 | 34 | 13 |
| Gudmund. | Nebraska: Hooker Co.,<br>Gudmundsen Sand Hills<br>laboratory, Whitman, NE | N42°04.881' | W101°28.154' | SH | 24 | 23 | 24 |
| SHGC | Nebraska: Hooker Co., Sand<br>Hills Golf Club, Mullen, NE | N41°51.605' | W101°05.117' | SH | 15 | 18 | 15 |
| Arapaho | Nebraska: Arthur Co.,<br>Arapaho Prairie, Arthur, NE | N41°28.939' | W101°50.966' | SH | 26 | 26 | 26 |
| Lemoyne | Nebraska: Keith Co.,<br>Lemoyne, NE | N41°17.098' | W101°50.329' | SH | 14 | 14 | 14 |
| Ogallala | Nebraska: Keith Co.,<br>cornfield in Ogallala, NE | N41°09.824' | W101°39.640' | Off | 24 | 24 | 24 |
| Grant | Nebraska: Perkins Co.,<br>cornfield in Grant, NE | N40°54.612' | W101°43.631' | Off | 30 | 30 | 30 |
| Imperial | Nebraska: Chase Co.,<br>cornfield in Imperial, NE | N40°32.270' | W101°37.240' | Off | 24 | 25 | 25 |

The name and location of each of 11 sampling sites are provided. “Habitat” refers to whether the site is on or off of the Sand Hills (“SH” and “Off”, respectively). “N” indicates the final sample sizes for analysis of phenotypic (“pheno”), environmental (“soil”), and genetic (“geno”) variation. Note: juveniles were genotyped but not scored for phenotype.

**Supplementary Table 2.** Strength of phenotypic association for previously identified candidate SNPs.

| Candidate (2013) | Traits (2013) | Position in scaffold | Dorsal brightness | Dorsal hue | Ventral brightness | D-V boundary | Tail stripe |
| --- | --- | --- | --- | --- | --- | --- | --- |
| 31327 | Tail stripe | 25175452 | **0.082** | 0.024 | 0.045 | **0.046** | **0.175** |
| 77047 | D-V boundary | 25128625 | 0.055 | 0.023 | 0.032 | 0.024 | 0.059 |
| 118834 | Dorsal hue | 25085120 | **0.203** | **0.047** | **0.122** | 0.026 | **0.123** |
| $\Delta_{\text{ser}}$ | Tail stripe; ventral color | 25075943 | **0.543** | **0.049** | **0.096** | **0.497** | **0.137** |
| 147347 | Tail stripe | 25055240 | 0.051 | 0.035 | 0.037 | 0.026 | 0.058 |
| 147789 | D-V boundary | 25054805 | **0.069** | 0.023 | 0.044 | 0.037 | 0.049 |
| Mean |  |  | 0.055 | 0.026 | 0.038 | 0.033 | 0.058 |

Candidate (2013) and Traits (2013) refer to significant genotype-phenotype associations identified in Linnen *et al.* (2013); remaining columns pertain to this study. Values in cells correspond to the posterior inclusion probabilities (PIP) for each trait and each candidate SNP (location on scaffold 16 is in base pairs). Gray boxes indicate SNPs that exceed our PIP threshold ( $>0.1$ ). Bolded numbers exceed background levels of association for that trait (mean + 0.01; mean PIP across all SNPs for each trait is given in the last row).

**Supplementary Table 3.** Hyper-parameters describing the genetic architecture of five color traits.

| Trait | h | PVE | rho | PGE | n_gamma |
| --- | --- | --- | --- | --- | --- |
| Dorsal brightness | 0.66 (0.37, 0.92) | 0.79 (0.60, 0.95) | 0.76 (0.43, 0.98) | 0.87 (0.68, 0.99) | 132.10 (7, 291) |
| Dorsal hue | 0.32 (0.10, 0.59) | 0.31 (0.11, 0.54) | 0.42 (0.03, 0.92) | 0.41 (0, 0.93) | 73.44 (0, 276) |
| Ventral brightness | 0.40 (0.20, 0.63) | 0.45 (0.25, 0.66) | 0.57 (0.18, 0.93) | 0.62 (0.23, 0.95) | 92.43 (3, 277) |
| D-V boundary | 0.57 (0.30, 0.82) | 0.68 (0.48, 0.87) | 0.78 (0.47, 0.98) | 0.87 (0.68, 0.99) | 62.37 (6, 214) |
| Tail stripe | 0.80 (0.53, 1.00) | 0.88 (0.71, 1.00) | 0.82 (0.60, 0.98) | 0.88 (0.73, 0.99) | 196.64 (60, 296) |

All parameters were estimated using a BSLMM model in GEMMA, 8,616 SNPs, and a relatedness matrix estimated from genome-wide SNPs. Parameter means and 95% credible intervals (lower bound, upper bound) were calculated from ten independent runs per trait. Interpretation of hyper-parameters is as follows: *PVE*, the total proportion of phenotypic variance that is explained by genotype; *PGE*, the proportion of genetic variance explained by sparse (*i.e.*, major) effects; *h*, an approximation used for prior specification of the expected value of *PVE*; *rho*, an approximation used for prior specification of the expected value of *PGE*; *n\_gamma*, the expected number of sparse (major) effect SNPs.

**Supplementary Table 4.** Posterior inclusion probabilities for candidate SNPs identified in association mapping analysis of all SNPs.

| SNP Position | Dorsal<br>brightness | Dorsal<br>hue | Ventral<br>brightness | D-V<br>boundary | Tail<br>stripe | Maximum |
| --- | --- | --- | --- | --- | --- | --- |
| scaffold00016:25175228 | 0.0470 | 0.0086 | 0.0279 | 0.7796 | 0.0412 | 0.7796 |
| scaffold00016:25075943 | 0.4683 | 0.0196 | 0.0479 | 0.2306 | 0.0818 | 0.4683 |
| NW_006502179.1:31437 | 0.0735 | 0.0073 | 0.0203 | 0.0036 | 0.3777 | 0.3777 |
| scaffold00016:25075903 | 0.0237 | 0.0080 | 0.0107 | 0.0350 | 0.3404 | 0.3404 |
| NW_006501090.1:4441929 | 0.0287 | 0.0077 | 0.0103 | 0.3168 | 0.0177 | 0.3168 |
| NW_006501041.1:4744196 | 0.0160 | 0.0084 | 0.0091 | 0.2539 | 0.0248 | 0.2539 |
| NW_006501185.1:901540 | 0.2517 | 0.0089 | 0.0121 | 0.0048 | 0.0214 | 0.2517 |
| scaffold00016:25076212 | 0.0428 | 0.0076 | 0.0227 | 0.0156 | 0.2495 | 0.2495 |
| NW_006501416.1:2319587 | 0.0435 | 0.0068 | 0.0249 | 0.0034 | 0.2466 | 0.2466 |
| scaffold00016:25085405 | 0.0111 | 0.0145 | 0.0109 | 0.0295 | 0.2251 | 0.2251 |
| NW_006501625.1:1977638 | 0.2184 | 0.0062 | 0.0072 | 0.0226 | 0.0137 | 0.2184 |
| NW_006501710.1:808272 | 0.0122 | 0.0102 | 0.0124 | 0.0178 | 0.2157 | 0.2157 |
| NW_006501970.1:52602 | 0.0122 | 0.0085 | 0.0104 | 0.2151 | 0.0535 | 0.2151 |
| scaffold00016:25074418 | 0.0153 | 0.0077 | 0.0091 | 0.0067 | 0.2151 | 0.2151 |
| scaffold00016:25173672 | 0.0735 | 0.0081 | 0.0925 | 0.0316 | 0.2147 | 0.2147 |
| NW_006501363.1:1300597 | 0.0149 | 0.0094 | 0.0088 | 0.0038 | 0.2106 | 0.2106 |
| NW_006501215.1:2183663 | 0.0109 | 0.0075 | 0.0118 | 0.0045 | 0.2096 | 0.2096 |
| NW_006501164.1:562237 | 0.0149 | 0.0077 | 0.0084 | 0.0040 | 0.2042 | 0.2042 |
| NW_006501505.1:1234861 | 0.0144 | 0.0082 | 0.0222 | 0.0036 | 0.2035 | 0.2035 |
| NW_006501064.1:4448524 | 0.0104 | 0.1988 | 0.0079 | 0.0039 | 0.0279 | 0.1988 |
| scaffold00016:25076355 | 0.0711 | 0.0072 | 0.0129 | 0.0121 | 0.1936 | 0.1936 |
| NW_006501066.1:6401068 | 0.0096 | 0.0070 | 0.0080 | 0.0044 | 0.1926 | 0.1926 |
| NW_006501363.1:1424874 | 0.0126 | 0.0099 | 0.0089 | 0.1915 | 0.0212 | 0.1915 |
| scaffold00016:25108248 | 0.1841 | 0.0073 | 0.0332 | 0.0077 | 0.0318 | 0.1841 |
| NW_006501066.1:6400593 | 0.0112 | 0.0092 | 0.0092 | 0.1833 | 0.0195 | 0.1833 |
| NW_006501258.1:1259668 | 0.0531 | 0.0076 | 0.0156 | 0.0633 | 0.1811 | 0.1811 |
| scaffold00016:25078686 | 0.1800 | 0.0070 | 0.0753 | 0.0152 | 0.0334 | 0.1800 |
| scaffold00016:25175452 | 0.0577 | 0.0073 | 0.0178 | 0.0167 | 0.1751 | 0.1751 |
| scaffold00016:25111157 | 0.1734 | 0.0079 | 0.0379 | 0.0133 | 0.0366 | 0.1734 |
| scaffold00016:25175879 | 0.0137 | 0.0097 | 0.0112 | 0.1706 | 0.0166 | 0.1706 |
| NW_006501149.1:6204758 | 0.0198 | 0.0083 | 0.0138 | 0.0088 | 0.1694 | 0.1694 |
| scaffold00016:25195324 | 0.0113 | 0.0084 | 0.0122 | 0.1643 | 0.0750 | 0.1643 |
| scaffold00016:25117322 | 0.0351 | 0.0072 | 0.0170 | 0.0041 | 0.1628 | 0.1628 |
| scaffold00016:25033802 | 0.0143 | 0.0090 | 0.0196 | 0.0050 | 0.1598 | 0.1598 |
| scaffold00016:25085108 | 0.1598 | 0.0210 | 0.0926 | 0.0050 | 0.0740 | 0.1598 |
| scaffold00016:25058748 | 0.0115 | 0.0119 | 0.0114 | 0.0089 | 0.1587 | 0.1587 |
| NW_006501112.1:2369721 | 0.0428 | 0.0071 | 0.1560 | 0.0034 | 0.0215 | 0.1560 |
| scaffold00016:25103837 | 0.0402 | 0.0141 | 0.0142 | 0.0092 | 0.1547 | 0.1547 |
| scaffold00016:25103836 | 0.0413 | 0.0143 | 0.0145 | 0.0094 | 0.1543 | 0.1543 |
| scaffold00016:25066132 | 0.0134 | 0.0077 | 0.0124 | 0.1537 | 0.0149 | 0.1537 |
| scaffold00016:25108191 | 0.1531 | 0.0078 | 0.0345 | 0.0074 | 0.0306 | 0.1531 |
| scaffold00016:25085120 | 0.1502 | 0.0226 | 0.0793 | 0.0055 | 0.0678 | 0.1502 |
| scaffold00016:25063343 | 0.0206 | 0.0080 | 0.0088 | 0.1491 | 0.0166 | 0.1491 |
| NW_006501047.1:4679252 | 0.0257 | 0.0092 | 0.0079 | 0.1488 | 0.0225 | 0.1488 |
| scaffold00016:25113309 | 0.1483 | 0.0109 | 0.0089 | 0.0058 | 0.0160 | 0.1483 |
| NW_006501634.1:1603622 | 0.0192 | 0.0076 | 0.0112 | 0.0042 | 0.1478 | 0.1478 |
| scaffold00016:25074446 | 0.0128 | 0.0167 | 0.0082 | 0.0062 | 0.1415 | 0.1415 |
| NW_006501392.1:259982 | 0.0163 | 0.0078 | 0.0080 | 0.0041 | 0.1403 | 0.1403 |
| scaffold00016:25076229 | 0.0242 | 0.0085 | 0.0160 | 0.0157 | 0.1320 | 0.1320 |
| scaffold00016:25039123 | 0.0120 | 0.0086 | 0.0103 | 0.1317 | 0.0182 | 0.1317 |
| NW_006501505.1:1621286 | 0.1308 | 0.0067 | 0.0116 | 0.0090 | 0.0157 | 0.1308 |
| NW_006501476.1:2057524 | 0.0274 | 0.0095 | 0.0098 | 0.0051 | 0.1298 | 0.1298 |
| NW_006501197.1:6731032 | 0.0129 | 0.0081 | 0.0092 | 0.0056 | 0.1281 | 0.1281 |
| NW_006501271.1:2908119 | 0.0296 | 0.0083 | 0.0095 | 0.0148 | 0.1253 | 0.1253 |
| NW_006501154.1:2893063 | 0.0111 | 0.0075 | 0.0074 | 0.0036 | 0.1249 | 0.1249 |
| scaffold00016:25066476 | 0.0108 | 0.0079 | 0.0068 | 0.1240 | 0.0151 | 0.1240 |
| NW_006501355.1:2485903 | 0.1238 | 0.0125 | 0.0161 | 0.0046 | 0.0198 | 0.1238 |
| NW_006501156.1:4189217 | 0.0103 | 0.0147 | 0.0076 | 0.0043 | 0.1227 | 0.1227 |
| scaffold00016:25098160 | 0.0145 | 0.0080 | 0.0117 | 0.1215 | 0.0349 | 0.1215 |
| scaffold00016:25063986 | 0.0182 | 0.0080 | 0.0104 | 0.1214 | 0.0383 | 0.1214 |
| NW_006502017.1:177456 | 0.1211 | 0.0106 | 0.0098 | 0.0037 | 0.0196 | 0.1211 |
| NW_006501451.1:1085407 | 0.0134 | 0.0082 | 0.0103 | 0.0124 | 0.1205 | 0.1205 |
| scaffold00016:25084934 | 0.1204 | 0.0118 | 0.0357 | 0.0087 | 0.0465 | 0.1204 |
| NW_006504112.1:718 | 0.0130 | 0.0083 | 0.0077 | 0.0040 | 0.1197 | 0.1197 |
| NW_006501124.1:2767575 | 0.0094 | 0.0082 | 0.0097 | 0.0043 | 0.1186 | 0.1186 |
| NW_006501363.1:1301032 | 0.0196 | 0.0079 | 0.0099 | 0.1168 | 0.0190 | 0.1168 |
| scaffold00016:25085230 | 0.0102 | 0.0069 | 0.0089 | 0.0038 | 0.1160 | 0.1160 |
| scaffold00016:25173030 | 0.0315 | 0.0086 | 0.0131 | 0.0109 | 0.1158 | 0.1158 |
| NW_006501178.1:430728 | 0.0144 | 0.0074 | 0.1152 | 0.0045 | 0.0171 | 0.1152 |
| NW_006501342.1:346813 | 0.0334 | 0.0079 | 0.0205 | 0.0153 | 0.1152 | 0.1152 |
| NW_006502126.1:565845 | 0.0105 | 0.0071 | 0.0075 | 0.0063 | 0.1150 | 0.1150 |
| NW_006501818.1:1377012 | 0.1143 | 0.0074 | 0.0142 | 0.0043 | 0.0213 | 0.1143 |
| NW_006501064.1:4448280 | 0.1132 | 0.0070 | 0.0071 | 0.0037 | 0.0262 | 0.1132 |
| scaffold00016:25086542 | 0.0306 | 0.0072 | 0.0321 | 0.0074 | 0.1127 | 0.1127 |
| scaffold00016:25075977 | 0.1126 | 0.0117 | 0.0368 | 0.0288 | 0.0160 | 0.1126 |
| NW_006501049.1:2210807 | 0.0131 | 0.0108 | 0.0076 | 0.0047 | 0.1122 | 0.1122 |
| NW_006502572.1:27506 | 0.0134 | 0.0094 | 0.0082 | 0.0561 | 0.1114 | 0.1114 |
| scaffold00016:25126621 | 0.0149 | 0.0232 | 0.0084 | 0.0047 | 0.1109 | 0.1109 |
| scaffold00016:25157370 | 0.0174 | 0.0083 | 0.0100 | 0.0057 | 0.1087 | 0.1087 |
| NW_006501094.1:2379444 | 0.0116 | 0.0080 | 0.0227 | 0.1080 | 0.0184 | 0.1080 |
| NW_006501818.1:1376978 | 0.1078 | 0.0076 | 0.0167 | 0.0048 | 0.0190 | 0.1078 |
| scaffold00016:25175932 | 0.0605 | 0.0086 | 0.0349 | 0.1065 | 0.0921 | 0.1065 |
| NW_006501066.1:6860433 | 0.0388 | 0.0295 | 0.0134 | 0.0051 | 0.1063 | 0.1063 |
| NW_006501280.1:3657919 | 0.1059 | 0.0073 | 0.0085 | 0.0067 | 0.0162 | 0.1059 |
| scaffold00016:25195321 | 0.0125 | 0.0097 | 0.0106 | 0.1053 | 0.0944 | 0.1053 |
| NW_006501131.1:1733529 | 0.0123 | 0.0103 | 0.0091 | 0.0044 | 0.1043 | 0.1043 |
| NW_006501237.1:5407969 | 0.0137 | 0.0095 | 0.0092 | 0.0070 | 0.1035 | 0.1035 |
| scaffold00016:25107989 | 0.1035 | 0.0084 | 0.0492 | 0.0151 | 0.0521 | 0.1035 |
| scaffold00016:25155955 | 0.0099 | 0.0071 | 0.0096 | 0.0050 | 0.1025 | 0.1025 |
| NW_006501392.1:260035 | 0.1024 | 0.0085 | 0.0109 | 0.0051 | 0.0194 | 0.1024 |
| scaffold00016:25064166 | 0.0142 | 0.0080 | 0.0102 | 0.1023 | 0.0239 | 0.1023 |
| NW_006501363.1:1420928 | 0.0088 | 0.0070 | 0.0085 | 0.1018 | 0.0137 | 0.1018 |
| NW_006501387.1:1348066 | 0.1016 | 0.0080 | 0.0568 | 0.0059 | 0.0211 | 0.1016 |
| scaffold00016:25063484 | 0.0136 | 0.0078 | 0.0152 | 0.1011 | 0.0282 | 0.1011 |
| NW_006501267.1:5399680 | 0.0315 | 0.0080 | 0.0095 | 0.0062 | 0.1011 | 0.1011 |

Position of each SNP is given as scaffold:position (in base pairs) in scaffold. *Agouti* is located on scaffold 16. All other scaffolds are “non-*Agouti*”. SNP posterior inclusion probabilities (PIPs) were estimated using a BSLMM model in GEMMA, 8,616 SNPs, and a relatedness matrix estimated from genome-wide SNPs. SNPs were included as candidates in this table if they had a PIP>0.10 for at least one pigmentation trait.

**Supplementary Table 5.**
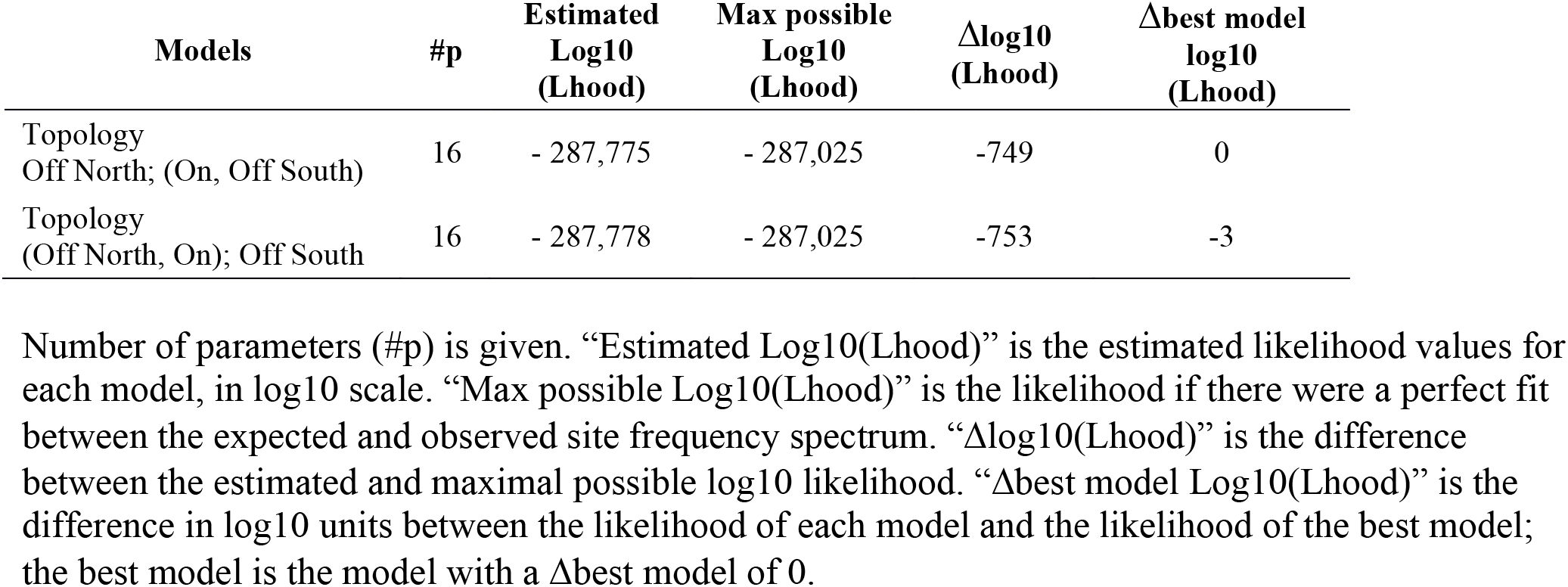
Likelihood values for each model, obtained with the full dataset including linked SNPs (3D-MSFS).

**Supplementary Table 6.** Parameter estimates obtained for the different demographic models.

| Parameters | Point estimates |  | 95% Confidence intervals |  |
| --- | --- | --- | --- | --- |
|  | Worst model<br>Topology<br>(Off N, On), Off S | Best model<br>Topology<br>Off N;<br>(On, Off S) | Best model<br>Topology<br>Off N;(On, Off S) |  |
| *Effective sizes (diploid Ne)* |  |  |  |  |
| N ancestral | 19,911 | 17,262 | 5,679 | 25,386 |
| N Off North | 38,343 | 39,847 | 27,890 | 58,658 |
| N On | 33,871 | 43,708 | 30,098 | 51,162 |
| N Off South | 60,166 | 54,568 | 39,936 | 68,150 |
| N ancNorth | 87,790 | 66,502 | 38,645 | 86,798 |
| N ancSouth | 55,416 | 56,768 | 30,953 | 84,090 |
| N BotOn (100 generations) | 32 | 17 | 6 | 209 |
| N BotAncestral (100 generations) | 13 | 884 | 18 | 1,379 |
| *Time of events (years)* |  |  |  |  |
| Time Split On | 4,026 | 3,709 | 3,400 | 7,942 |
| Time Split ancestral | 39,410 | 45,471 | 37,073 | 57,782 |
| *Scaled migration rates (2Nm, number of immigrants per generation)* |  |  |  |  |
| On to Off N | 13.93 | 12.46 | 2.6 | 18.2 |
| Off N to On | 6.97 | 6.40 | 1.4 | 13.1 |
| Off S to On | 17.82 | 18.28 | 12.5 | 24.0 |
| On to Off S | 0.37 | 3.62 | 0.0 | 11.3 |
| Between Off S and Off N | 4.99 | 4.87 | 1.9 | 8.6 |
| Between Off N (or Off S) and ancestral On-Off S (or On-Off N) | 5.6e-4 | 3.61e-4 | 0.0 | 0.3 |
| *Migration rates (m, proportion of migrants per generation)* |  |  |  |  |
| On to Off N | 1.59e-04 | 1.56e-04 | 2.36e-05 | 3.27e-04 |
| Off N to On | 9.03e-05 | 7.32e-05 | 1.79e-05 | 1.99e-04 |
| Off S to On | 2.31e-04 | 2.09e-04 | 1.54e-04 | 3.35e-04 |
| On to Off S | 2.67e-06 | 3.32e-05 | 3.24e-10 | 1.39e-04 |
| Between Off S and Off N | 5.72e-05 | 6.11e-05 | 3.42e-05 | 8.08e-05 |
| Between Off N (or Off S) and ancestral On-Off S (or On-Off N) | 2.84e-09 | 2.71e-09 | 6.64e-11 | 2.57e-06 |

**Supplementary Table 7.** *F_ST_* values in the *Agouti* locus.

| **Position** | **$F_{ST}$: On vs Off North** | **$F_{ST}$: On vs Off South** |
| --- | --- | --- |
| 25103904 | 0.442782 | 0.455174 |
| 25107989 | 0.475032 | 0.466737 |
| 25108113 | 0.452818 | 0.454937 |
| 25108191 | 0.451366 | 0.474141 |
| 25108248 | 0.451366 | 0.463734 |
| 25108273 | 0.422605 | 0.475471 |
| 25111157 | 0.457267 | 0.490538 |
| 25111390 | 0.402897 | 0.49633 |

Summary of $F_{ST}$ values “On” the Sand Hills relative to “Off North” of the Sand Hills and separately to “Off South” of the Sand Hills. SNP positions given in basepairs on scaffold 16.

**Supplementary Table 8.** Summary of the filtered datasets used for analysis.

|  | Population structure<br>(1 SNP per block) | Demographic<br>analyses | Association<br>mapping |
| --- | --- | --- | --- |
| Length blocks (kb) | 1.5 | 1.5 | 1 |
| Min. #accessible sites per block | 150 | 150 | NA |
| Min. median distance among SNPs within<br>a block (in bp) | 3 | 3 | NA |
| Min. depth of coverage DP | 4 | 4 | 2 |
| Min. individual mean DP | 8 | 8 | 2 |
| Max. DP | $2*mDP$ | $2*mDP$ | $2*mDP$ |
| Max. missing data per site allowed | 0.1 | 0 | 0.1 |
| Max. missing data per individual allowed | 0.25 | 0 | 0.25 |
| Minor allele frequency filter | $1/(2*n)$ | 0 | 0.01 |
| Number of individuals | 161 | 30 | 236 |
| Number of SNPs | 5412 | 140358 | 12920 |
| Number of accessible sites | NA | 2674174 | NA |

$mDP$ corresponds to the mean depth of coverage (DP) per individual, and $n$ corresponds to the number of individuals. Three columns provide details for each analyses presented – the dataset used to assess population structure, demographic analyses, and association mapping.

**Supplementary Table 9.** Summary of the dataset based on the random background genomic regions.

|  | #Blocks | #SNPs | #Accessible sites | Median<br>#SNPs/block | Median<br>#CallableSites/block |
| --- | --- | --- | --- | --- | --- |
| All blocks | 52566 | 649300 | 6224355 | 4 | 100 |
| Filtered blocks | 11770 | 284287 | 2814532 | 19 | 207 |

**Supplementary Table 10.** Inference of demographic history.

| Parameter tag | Range units | Parameter range |  |  |
| --- | --- | --- | --- | --- |
|  |  | lower | upper |  |
| N ancestral | uniform | 10000 | 1000000 | * |
| Time Split On | uniform | 200 | 50000 |  |
| Time Split ancestral | uniform | 1 | 50000 | <sup>a</sup> |
| N North | uniform | 1000 | 400000 |  |
| N On | uniform | 1000 | 400000 |  |
| N South | uniform | 1000 | 400000 |  |
| N ancNorth | uniform | 1000 | 400000 |  |
| N ancSouth | uniform | 1000 | 400000 |  |
| N BotOn | uniform | 10 | 3000 | * |
| 2Nm On to Off N | logunif | 1.00E-06 | 10 | * |
| 2Nm Off N to On | logunif | 1.00E-06 | 10 | * |
| 2Nm Off S to On | logunif | 1.00E-06 | 10 | * |
| 2Nm On to Off S | logunif | 1.00E-06 | 10 | * |
| 2Nm Off S to Off N | logunif | 1.00E-06 | 10 | * |
| 2Nm Off N (or Off S) to<br>ancestral On-Off S (or On-Off N) | logunif | 1.00E-06 | 10 | * |

**Supplementary Table 11.** Nested ANOVA

|  |  | Source | DF | SS | MS | F | P | PVE |
| --- | --- | --- | --- | --- | --- | --- | --- | --- |
| **<u>Environment</u>** | **Soil brightness** | Habitat | 1 | 129.66 | 129.66 | 307.40 | **<2e-16** | 0.48 |
|  |  | Habitat:Site | 9 | 30.37 | 3.37 | 8.00 | **1.89E-10** | 0.11 |
|  |  | Residuals | 260 | 109.68 | 0.42 |  |  | 0.41 |
| **<u>Color traits</u>** | **PC1 (dorsal bright.)** | Habitat | 1 | 87.01 | 87.01 | 218.56 | **<2e-16** | 0.39 |
|  |  | Habitat:Site | 9 | 37.68 | 4.19 | 10.52 | **8.07E-14** | 0.17 |
|  |  | Residuals | 254 | 101.12 | 0.40 |  |  | 0.45 |
|  | **PC2 (dorsal hue)** | Habitat | 1 | 17.60 | 17.60 | 18.41 | **2.53E-05** | 0.06 |
|  |  | Habitat:Site | 9 | 28.86 | 3.21 | 3.35 | **0.000671** | 0.10 |
|  |  | Residuals | 254 | 242.83 | 0.96 |  |  | 0.84 |
|  | **PC3 (ventral bright.)** | Habitat | 1 | 40.76 | 40.76 | 60.95 | **1.53E-13** | 0.15 |
|  |  | Habitat:Site | 9 | 62.16 | 6.91 | 10.33 | **1.42E-13** | 0.23 |
|  |  | Residuals | 254 | 169.85 | 0.67 |  |  | 0.62 |
|  | **PC4 (ventral hue)** | Habitat | 1 | 0.90 | 0.90 | 1.08 | 0.3 | 0.00 |
|  |  | Habitat:Site | 9 | 8.91 | 0.99 | 1.19 | 0.304 | 0.04 |
|  |  | Residuals | 254 | 212.12 | 0.84 |  |  | 0.96 |
|  | **D-V boundary** | Habitat | 1 | 59.32 | 59.32 | 104.62 | **<2e-16** | 0.25 |
|  |  | Habitat:Site | 9 | 33.93 | 3.77 | 6.65 | **1.54E-08** | 0.14 |
|  |  | Residuals | 255 | 144.60 | 0.57 |  |  | 0.61 |
|  | **Tail stripe** | Habitat | 1 | 80.65 | 80.65 | 170.21 | **<2e-16** | 0.35 |
|  |  | Habitat:Site | 9 | 29.34 | 3.26 | 6.88 | **7.54E-09** | 0.13 |
|  |  | Residuals | 251 | 118.94 | 0.47 |  |  | 0.52 |
| **<u>Non-color traits</u>** | **Total length** | Habitat | 1 | 0.09 | 0.09 | 0.11 | 0.74 | 0.00 |
|  |  | Habitat:Site | 9 | 60.27 | 6.70 | 8.54 | **3.89E-11** | 0.24 |
|  |  | Residuals | 250 | 196.08 | 0.78 |  |  | 0.76 |
|  | **Tail length** | Habitat | 1 | 0.14 | 0.14 | 0.18 | 0.672 | 0.00 |
|  |  | Habitat:Site | 9 | 49.12 | 5.46 | 7.19 | **2.85E-09** | 0.21 |
|  |  | Residuals | 250 | 189.91 | 0.76 |  |  | 0.79 |
|  | **Hind foot length** | Habitat | 1 | 2.10 | 2.10 | 3.01 | 0.084 | 0.01 |
|  |  | Habitat:Site | 9 | 105.40 | 11.71 | 16.81 | **<2e-16** | 0.38 |
|  |  | Residuals | 248 | 172.70 | 0.70 |  |  | 0.62 |
|  | **Ear length** | Habitat | 1 | 0.35 | 0.35 | 0.48 | 0.491 | 0.00 |
|  |  | Habitat:Site | 9 | 44.39 | 4.93 | 6.70 | **1.38E-08** | 0.20 |
|  |  | Residuals | 247 | 181.78 | 0.74 |  |  | 0.80 |

Nested ANOVA tables for environmental, color trait, and non-color trait measurements, with degrees of freedom (DF), sum of squares (SS), mean sum of squares (MS), *F*-statistic (F), *P*-value (P), and proportion variance explained (PVE). “Habitat” refers to whether samples were collected on or off the Sand Hills; “Habitat:Site” refers to collecting locations nested within habitat type. Significant *P*-values ( $\alpha = 0.05$ ) are bolded.

**Supplementary Figure 1.**
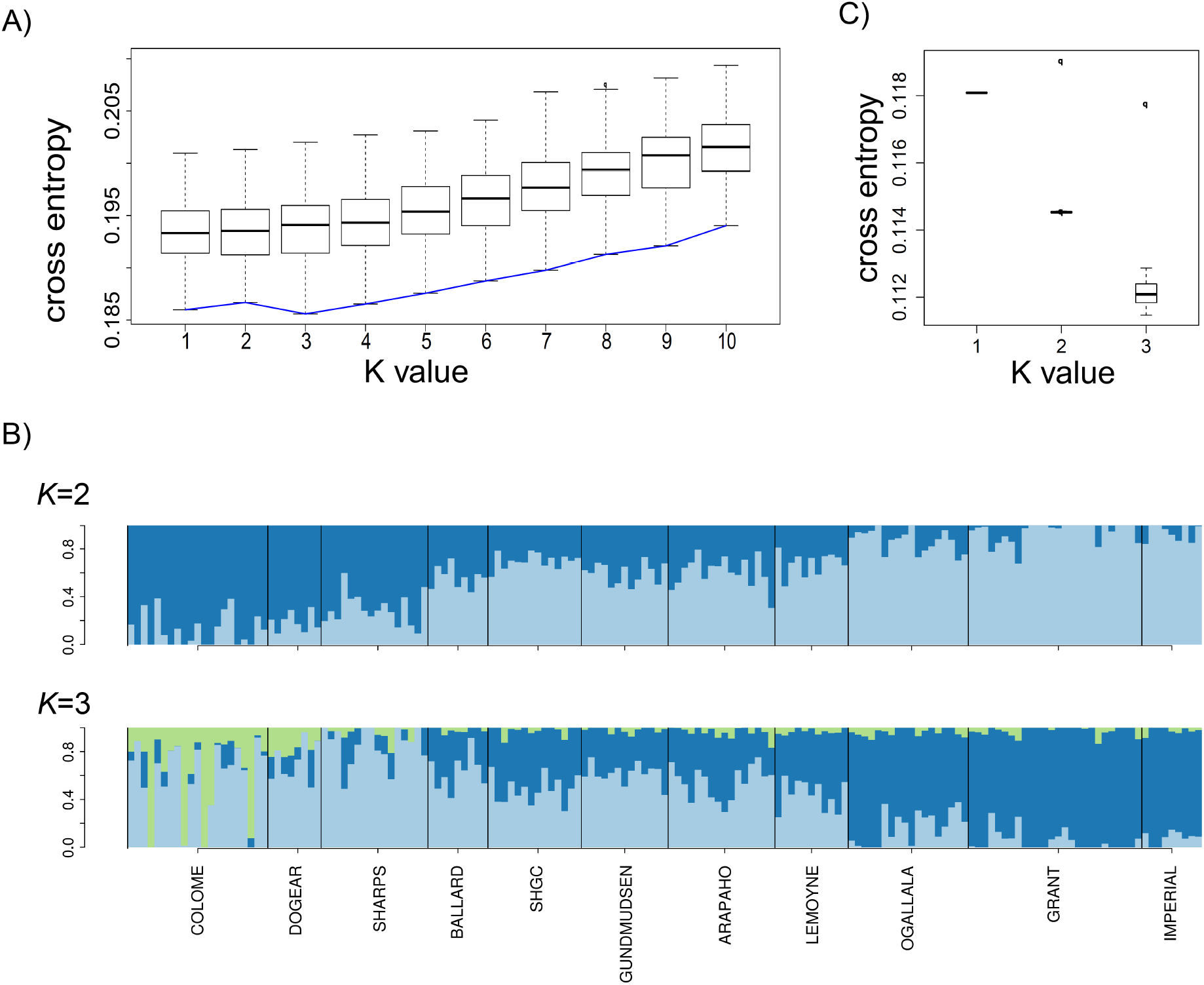
Population structure analyses with sNMF and TESS3. A) Crossentropy for each *K*-value. Boxplots correspond to the distribution of cross-entropy values obtained by performing 100 runs for each *K* value. Blue line represents the minimum cross-entropy for each *K* value. The distribution of cross-entropy is similar for values *K*=1−4, but the lowest *K* value (corresponding to the minimal cross-entropy) is *K*=3. B) Estimated ancestry proportions for each individual for *K*=2 and *K*=3 with sNMF, without accounting for geographic location of individuals. C) Cross-entropy for *K* values from 1 to 3 using the method accounting for geographic information implemented in TESS3. Boxplots correspond to the distribution of cross-entropy values obtained by performing 1000 runs for each *K* value. The blue line represents the minimum cross-entropy for each *K* value. When accounting for the geographic sampling location of individuals, the cross-entropy decreases for *K* from 1 to 3, with a minimum value at 3. Results reported with lambda value of 0.01, but similar results were obtained when changing the lambda values from 0.01 to 0.05 (values nearer to zero give less weight to the geographic location of individuals).

**Supplementary Figure 2.**
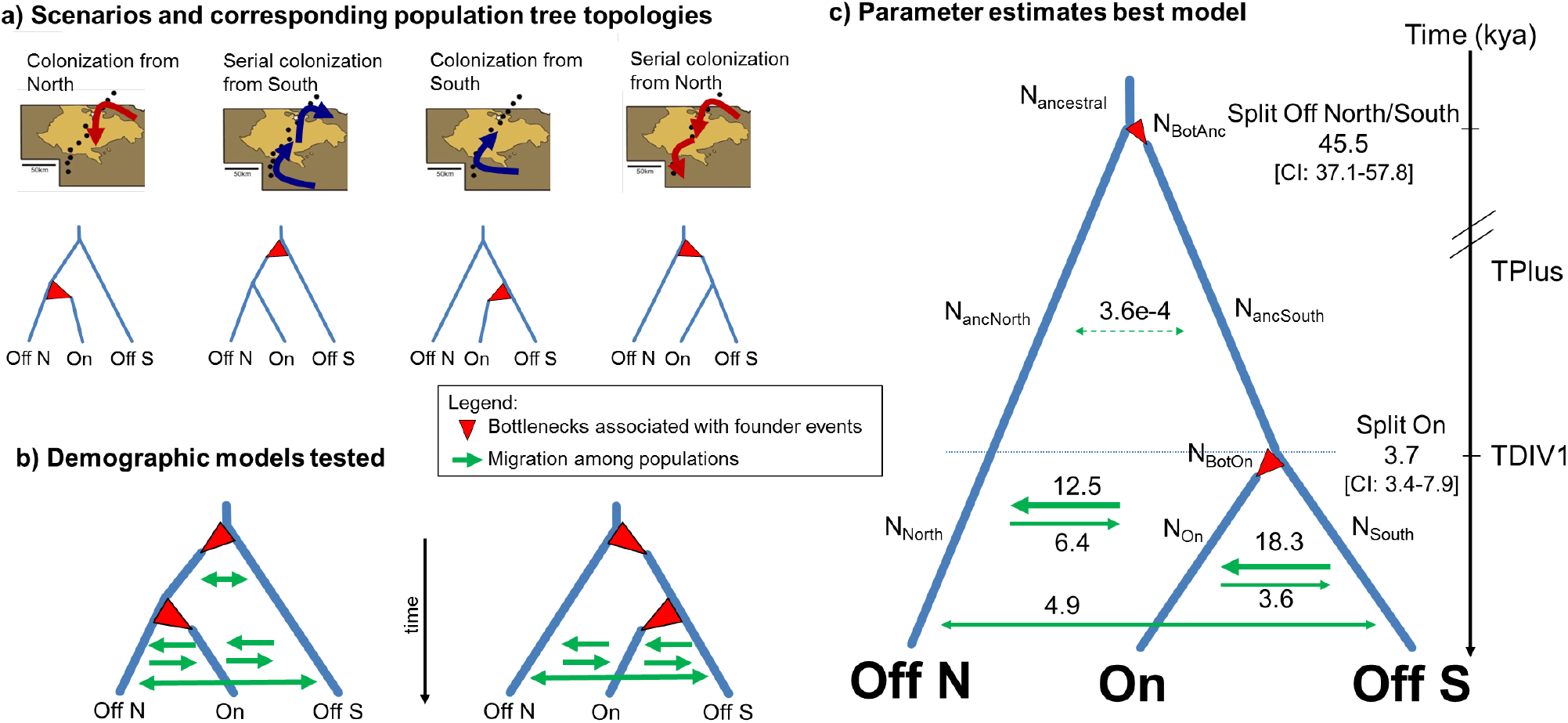
Demographic modeling favors colonization from the south. (**a**) Four scenarios were tested and the corresponding three-population model tree topologies are provided. The three samples correspond to populations north of the Sand Hills (Off N), on the Sand Hills (On), and south of the Sand Hills (Off S). Although alternative scenarios might have identical population trees, they can be distinguished as founder events associated with the colonization of the Sand Hills are expected to occur on different lineages. (**b**) Demographic models tested with two topologies, allowing for up to two bottlenecks associated with the population split, and gene flow among populations. Under this modeling framework, we can distinguish among scenarios with the same tree topology by inferring the strength of the bottlenecks associated with the split of the Sand Hills populations (indicated as red arrows), representing potential founder events related with the colonization of the Sand Hills. For instance, for the tree topology ((Off N, On), Off S), if inferred parameters indicate a stronger bottleneck associated with the Sand Hills (“On”), it would support a colonization from the north scenario, whereas a stronger bottleneck associated with the split of the ancestor of “Off N” and “On” would suggest a serial colonization from the south. (**c**) Parameter estimates under the best model, supporting a southern colonization route. Parameter estimates for scaled immigration rates *2Nm* (*i.e.*, average number of immigrants per generation) are shown above or below corresponding arrows, and the time of events (given in thousand years (kya)) is indicated at the nodes, assuming 2.5 generations per year and a mutation rate of 3.67×10^−8^ per site per generation. Parameter tags are indicated next to each corresponding parameter. Demographic modeling was based on the threedimensional minor allele frequency spectrum (3D MSFS), considering models with three populations. The 3D folded SFS was generated by dividing the data set into blocks of 1.5kb, sampling ten individuals for each block from each population without missing data – resulting in a 3D folded SFS with 140,358 SNPs.

**Supplementary Figure 3.**
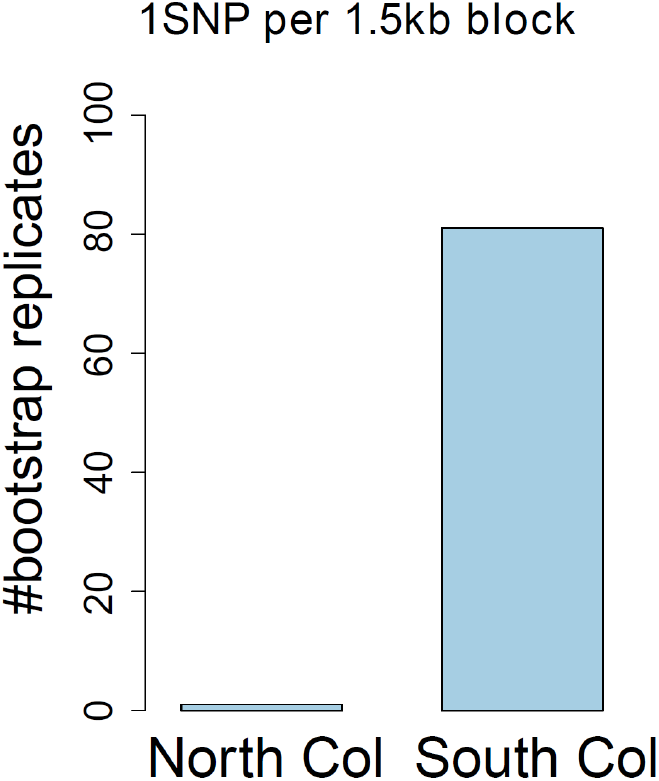
Model support based on Akaike’s information criterion (AIC) across bootstrap replicates. Only SNPs further than 1.5kb apart were considered, and hence each bootstrap replicate was obtained by sampling one SNP per 1.5kb block. Likelihoods were computed for each bootstrap replicate based on the expected site frequency spectrum for each model (approximated with 10×10^6^ coalescent simulations) using the parameter estimates that maximized the likelihood with a larger data set (140,358 SNPs), potentially including linked sites. The y-axis represents the number of bootstrap replicates supporting a given model. Each bootstrap replicate was assigned to the model with a relative likelihood (based on the AIC) larger than 0.95. On the y-axis, a value of 100 indicates that a given model was supported by all bootstrap replicates. Note: 14 bootstrap replicates could not be assigned to any model as the relative likelihood was below 0.95 for both models.

**Supplementary Figure 4:**
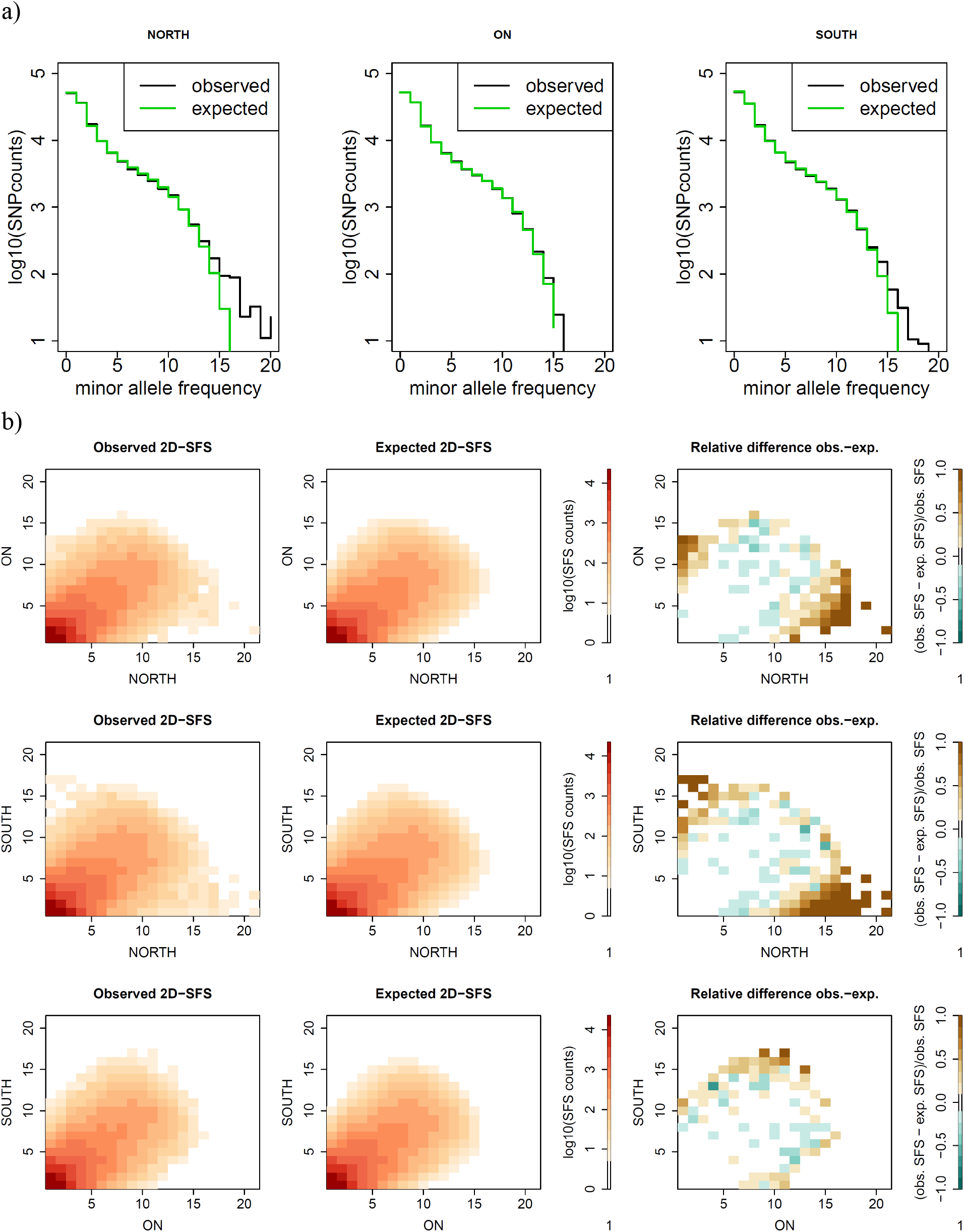
Observed marginal site frequency spectrum (SFS). (**a**) 1D and (**b**) 2D SFS and fit of the marginal minor allele frequency (MAF) spectrum for the best model. The south colonization model fits the MAF spectrum for each sample well. In panel b, the x-and y-axis represent the MAF. As can be seen by the marginal 2D observed SFS, the sample “On” the Sand Hills has more SNPs along the diagonal with the sample south “Off” the Sand Hills, suggesting it is more closely related to the southern population.

**Supplementary Figure 5.**
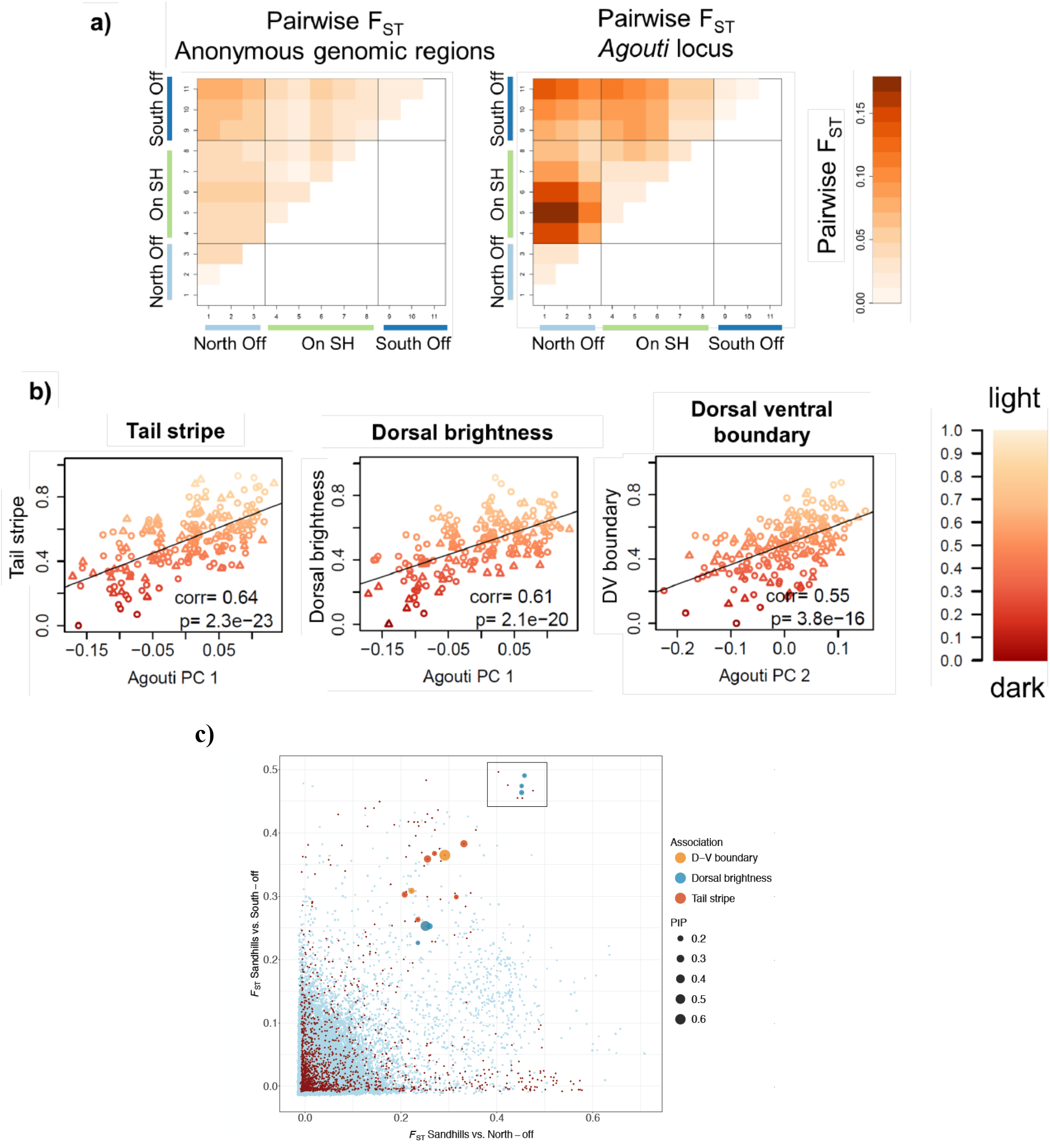
Comparison of genetic differentiation for random background genomic regions and *Agouti*. (**a**) *F_ST_* for samples along the transect. At the *Agouti* locus, higher *F_ST_* values are observed, particularly for comparisons between samples On (green) and Off (blue) of the Sand Hills. (**b**) Genetic differentiation at *Agouti* is correlated with phenotypic trait values. PC1 correlates with tail stripe and dorsal brightness, whereas PC2 correlates with the dorsal ventral boundary, in agreement with Linnen *et al*. 2013 (color-coded with darker phenotypes in darker shades). Circles represent males, and triangles females. (**c**) The x-axis gives *FST* values for On vs. North, and the y-axis gives values for On vs. South. Points highlighted by a square correspond to the highest peak of differentiation. For SNPs with *F_ST_* values above 0.2, their estimated contribution to the phenotypes considered in the association study are indicated by dot size (PIP > 0.15).

**Supplementary Figure 6.**
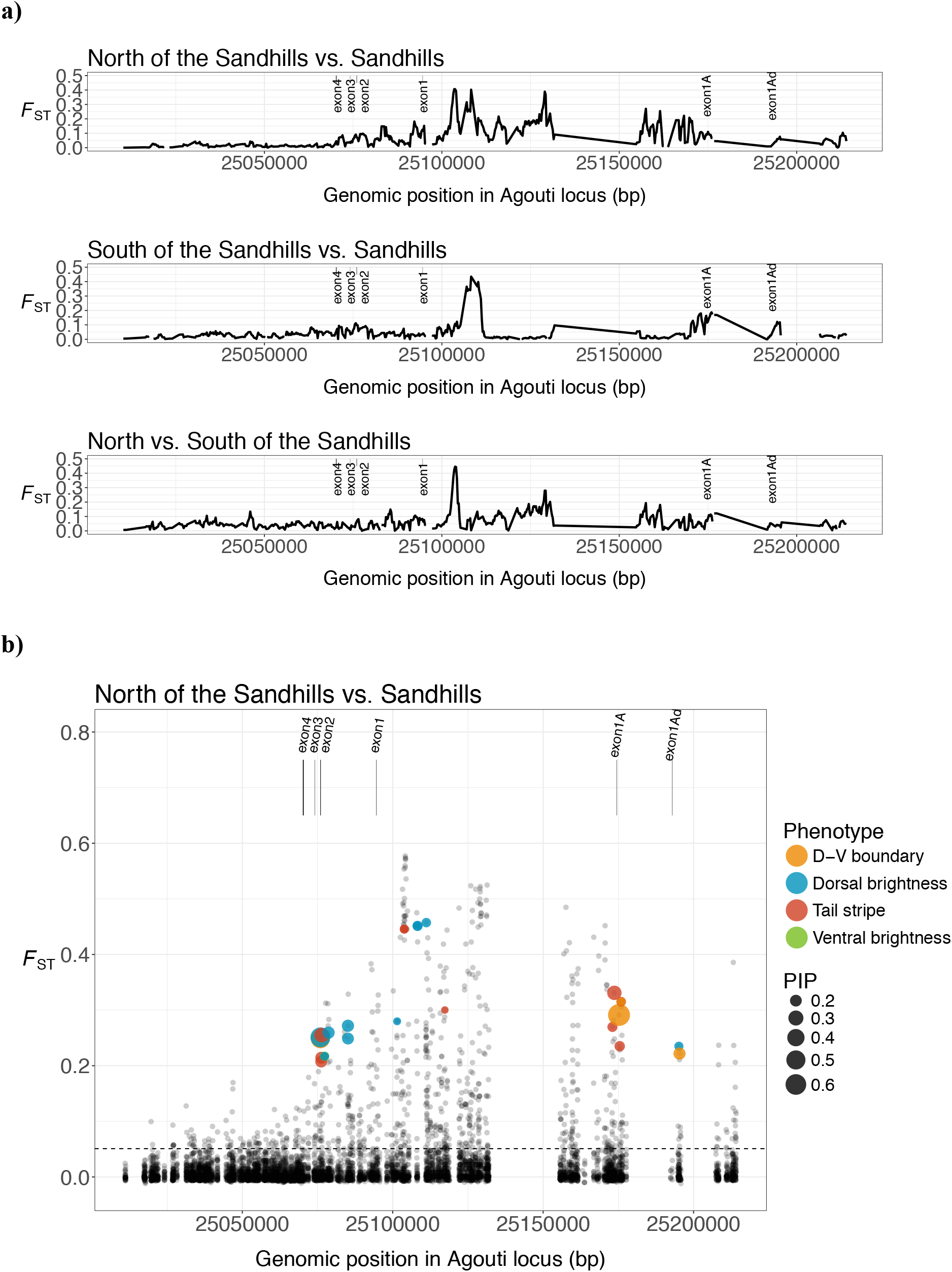
Genetic differentiation (*F_ST_*) between populations On and Off of the Sand Hills at *Agouti*. Values are calculated (**a**) in sliding windows (window size: 1kb, step size: 100bp) and (**b**) per SNP using VCFtools v.0.1.12b. Vertical lines correspond to the locations of known exons in the *Agouti* locus.

**Supplementary Figure 7.**
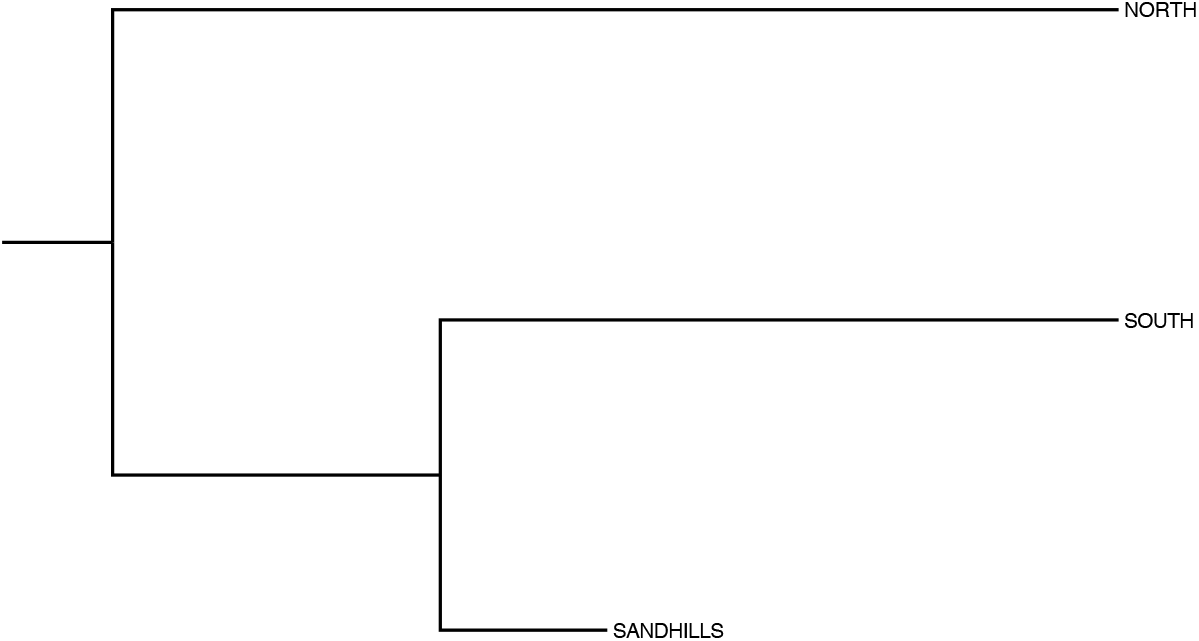
Graphical representation of the kinship matrix. This tree is output produced by HapFLK, using all background genomic regions (excluding *Agouti*). Note shorter branch lengths represent more similar allele frequencies among populations.

**Supplementary Figure 8.**
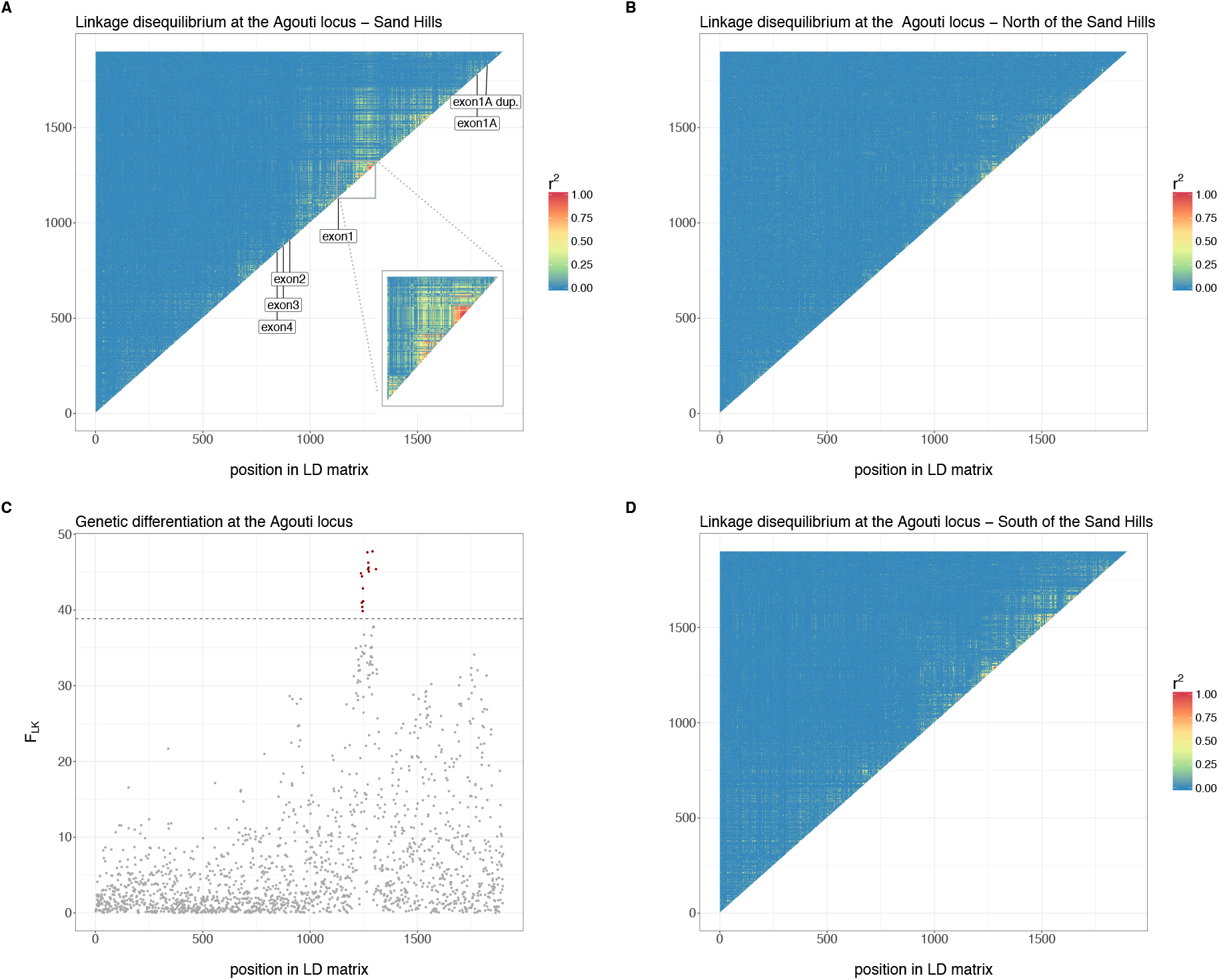
Patterns of linkage disequilibrium (LD) across the *Agouti* locus. *r^2^* values have been calculated with vcftools (--hap-r2) excluding all variants with a minor allele frequency strictly lower than 5%. The x-axis of the 4 panels are aligned. A) LD patterns in mice sampled at the Sand Hills. The inset highlights the fine-grained LD patterns around the significant peak of genetic differentiation (panel C). Approximate positions of all exons are indicated. B) LD patterns in mice sampled North of the Sand Hills. C) The FLK statistic represents variant-specific allelic differentiation across the 3 main geographic localities. The same filtering as in the LD analysis has been applied such that x-axes could be aligned. D) LD patterns in mice sampled South of the Sand Hills. For panels C and D, only variants present in the Sand Hills were used so that all three heat maps are constructed with an identical set of variants.

**Supplementary Figure 9.**
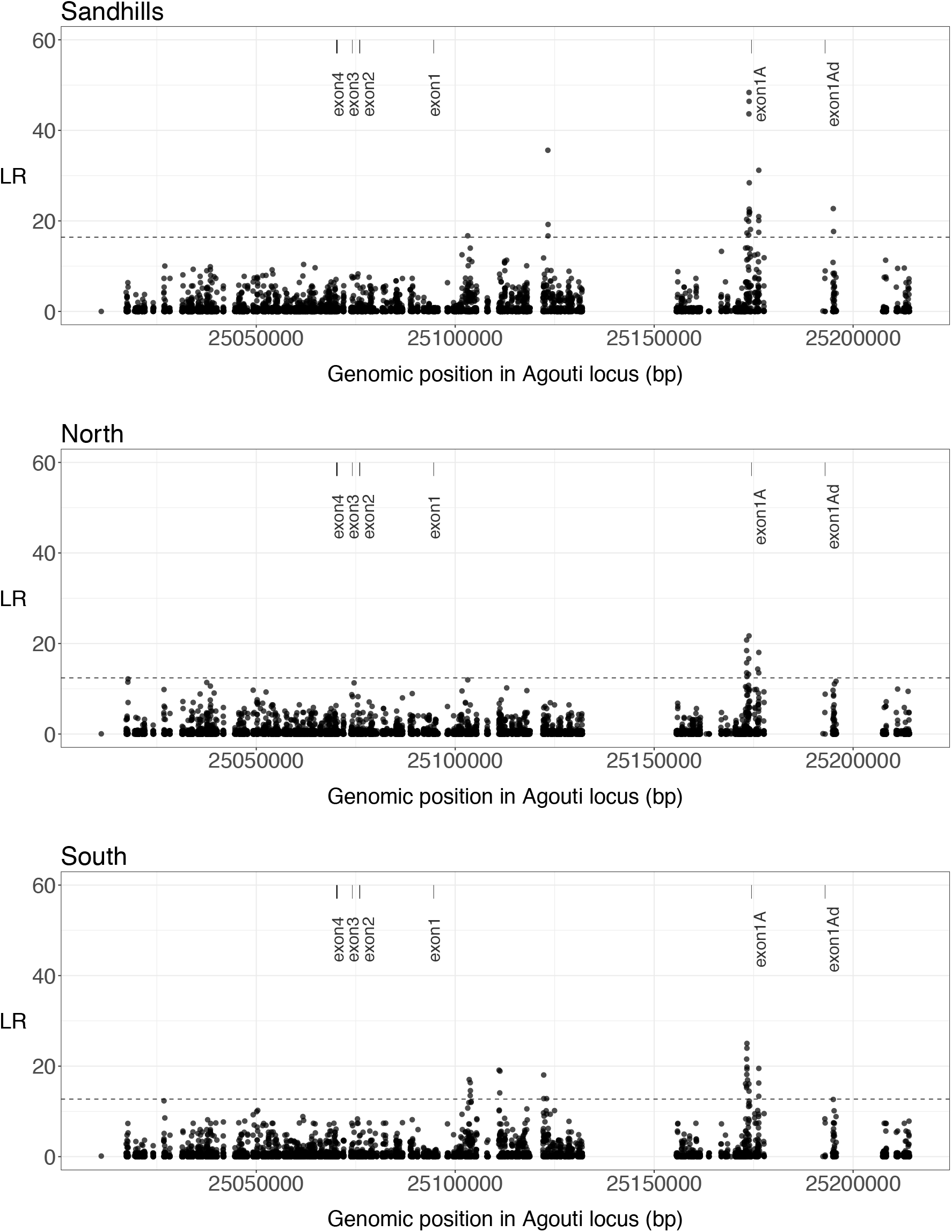
The composite likelihood ratio (CLR) test across the *Agouti* locus. The CLR test identifies SNPs that deviate from neutral expectations. The CLR statistic was obtained using a pre-computed SFS obtained from the background dataset for the (**a**) Sand Hills, (**b**) North of the Sand Hills and (**c**) South of the Sand Hills populations. Dotted horizontal lines correspond to a theoretical significance threshold calculated for the Sand Hills region using neutral simulations.

**Supplementary Figure 10.**
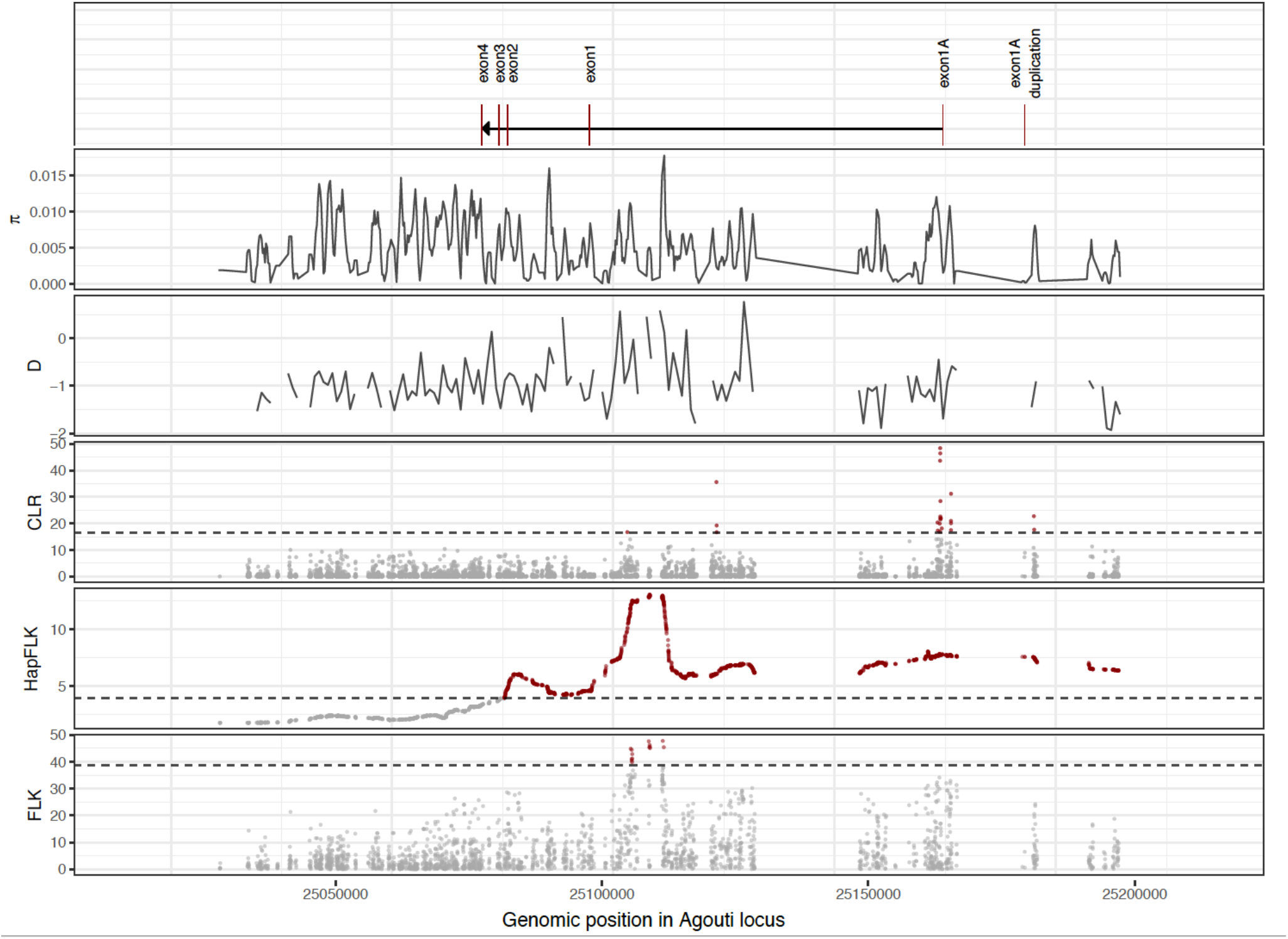
Comparison of tests of neutrality and selection across the Agouti locus. From the top, the exon structure, pairwise differences, Tajima’s D, CLR test, HapFLK, and FLK. The dashed lines indicate the significance threshold using parametric bootstrapping of the best-fitting demographic model.

**Supplemental Figure 11.**
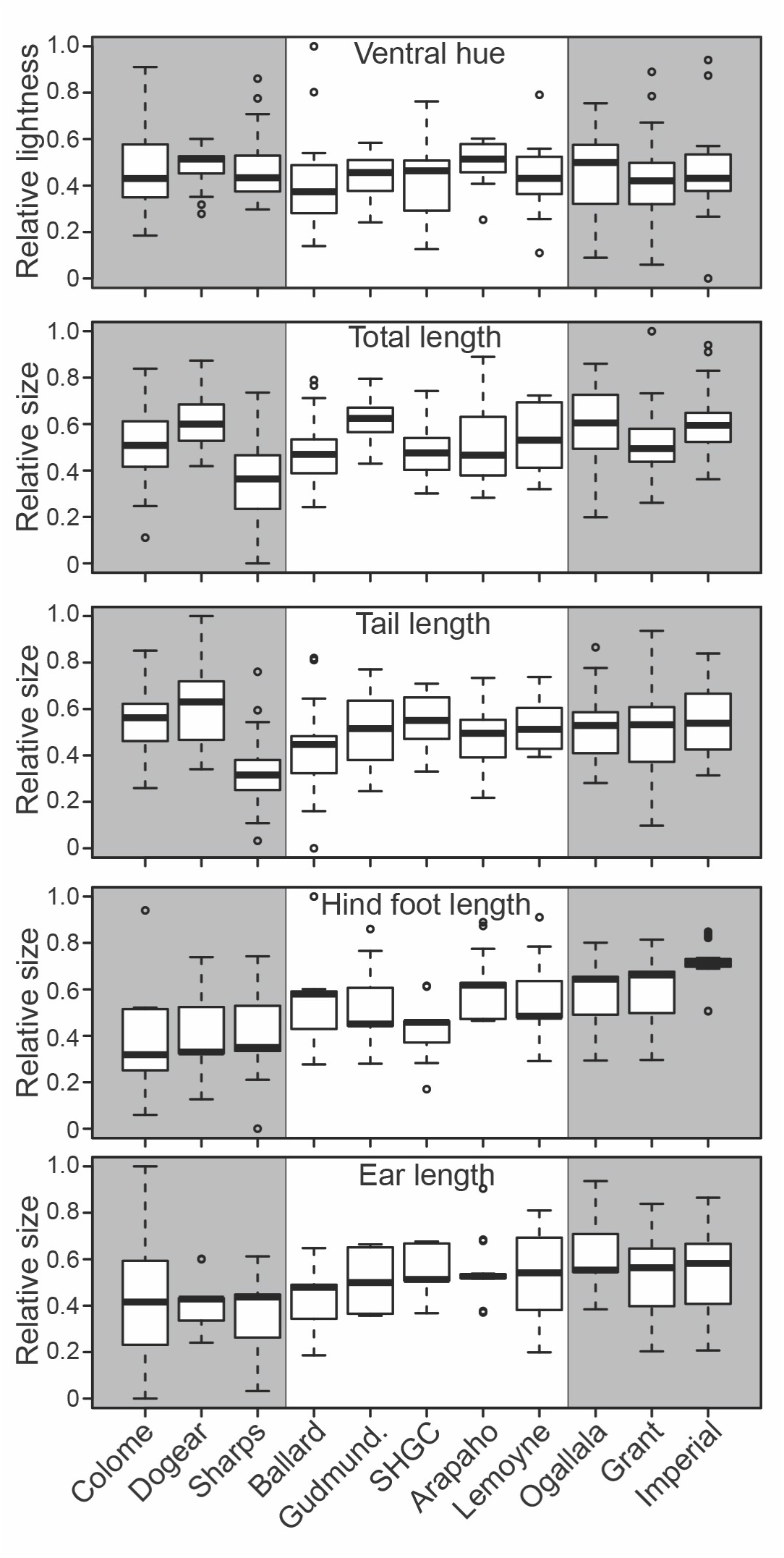
Ventral hue and non-color traits are not significantly different on and off the Sand Hills. Box plots for each sampling site were produced using scaled (range: 0-1), normal-quantile transformed values for: ventral hue (PC4), total body length, tail length, hind foot length, and ear length. In all plots, higher values correspond to lighter/brighter color or longer lengths. Grey shading indicates non-Sand Hills sites. Ventral hue did not differ significantly between habitats (on vs. off Sand Hills) or among sites (*P* > 0.05). All four length traits differed significantly among sites nested within habitats, but not among habitats.

